# Antimetastatic Sulfonate-Functionalized Mesoporous Silica Nanoparticles Enhance Irinotecan Stability and Delivery for Colorectal Cancer Treatment

**DOI:** 10.1101/2025.11.25.690433

**Authors:** Zih-An Chen, Cheng-Hsun Wu, Cong Kai Lin, Rong-Lin Zhang, Ting Chung Sun, Dai-Wen Li, Chung-Yuan Mou, Peilin Chen, Si-Han Wu, Yi-Ping Chen

## Abstract

Metastatic colorectal cancer (mCRC) remains a leading cause of cancer-related mortality, with Irinotecan (IRI) serving as a backbone chemotherapeutic despite its dose-limiting toxicities, instability of the active lactone form, and lack of intrinsic antimetastatic activity. To overcome these barriers, we developed sulfonate-functionalized mesoporous silica nanoparticles (MSNs), referred to as (SO_3_^-^)-MSN-PEG/TA, as a multifunctional nanocarrier for IRI delivery. This nanoformulation markedly enhanced drug loading efficiency and preserved over 90% of the lactone form for up to six months, enabling sustained release and improved pharmacological stability. In vitro studies demonstrated superior cellular uptake, enhanced apoptosis, and a reduced IC_50_ compared to free IRI. Beyond drug delivery, (SO_3_^-^)-MSN-PEG/TA exhibited intrinsic antimetastatic activity by modulating focal adhesion kinase (FAK)/paxillin signaling, thereby impairing cell migration and suppressing angiogenesis, along with efficient tumor accumulation through the enhanced permeability and retention (EPR) effect. Pharmacokinetic analysis further revealed that IRI@(SO_3_^-^)-MSN-PEG/TA prolonged systemic retention, maintaining higher IRI plasma concentrations compared with free IRI. IRI@(SO_3_^-^)-MSN-PEG/TA significantly inhibited both primary tumor growth and metastatic dissemination in orthotopic and heterotopic colorectal cancer models, while markedly reducing systemic toxicities and preserving bone marrow cellularity relative to free IRI and liposomal IRI (Onivyde). Collectively, this dual-functional nanomedicine provides an innovative therapeutic strategy that not only augments IRI efficacy but also confers metastasis suppression and favorable pharmacokinetics, addressing critical unmet needs in mCRC treatment. These findings highlight the translational potential of IRI@(SO_3_^-^)-MSN-PEG/TA as a safer and more effective therapy for mCRC.

## Introduction

Metastatic colorectal cancer (mCRC) remains a major global health challenge, ranking as the third most common cancer worldwide and a leading cause of cancer-related mortality.[1, 2] Despite advances in early detection and treatment, the prognosis for mCRC remains poor due to its aggressive progression and the high prevalence of metastatic relapse. Alarmingly, the incidence of mCRC in young adults under 50 years of age has been rising, further underscoring its clinical urgency.[3] Metastatic dissemination, which accounts for over 90% of cancer-associated deaths, highlights the need for therapeutic strategies that can simultaneously address primary tumor growth and metastasis spread.[4, 5]

Irinotecan (IRI) is a cornerstone chemotherapeutic for mCRC, typically administered as part of FOLFIRI regimens.[6] As a prodrug, IRI must be converted into its active metabolite, SN-38, through hydrolysis by carboxylesterases. SN-38 exerts cytotoxicity relies on inhibiting topoisomerase I and disrupting DNA replication in cancer cells.[7–9] However, the therapeutic benefit of IRI is limited by two critical challenges: (1) a pH-dependent interconversion between its active lactone and inactive carboxylate forms, where acidic conditions favor the active lactone form but physiological pH promotes conversion to the inactive carboxylate,[10, 11] and (2) severe dose-limiting toxicities, particularly neutropenia and diarrhea.[6, 12] Furthermore, the clinical efficacy of IRI is compromised by the drug resistance mechanisms in cancer cells, restricting its long-term effectiveness.

Nanomedicine has emerged as a promising strategy to overcome these challenges. In mCRC treatment, nanoparticles can deliver drugs such as IRI more directly to tumors, thereby increasing local drug concentrations and reducing systemic toxicity.[13–16] One representative example is liposomal IRI (Onivyde, MM-398), which was approved by the FDA in 2015 for the treatment of metastatic pancreatic ductal adenocarcinoma. It is administered in combination with fluorouracil and leucovorin for patients who have progressed following gemcitabine-based therapy, and more recently as part of the NALIRIFOX regimen (fluorouracil, leucovorin, and oxaliplatin) as first-line treatment at a dose of 50–70 mg/m² every two weeks. This liposomal formulation improves pharmacokinetics, prolongs systemic circulation, and enables sustained release of SN-38, leading to an overall survival benefit of approximately two months compared with fluorouracil and leucovorin alone.[17–19] However, Onivyde failed to demonstrate clinical benefit in mCRC patients during a phase II trial, likely due to the much higher dosage required for mCRC treatment (350 mg/m²) compared with pancreatic cancer, which resulted in intolerable adverse effects.[20] In addition, its relatively large particle size (∼110 nm) may have limited deep tumor penetration within the complex tumor matrix[21]. Importantly, Onivyde and most nanocarrier systems lack intrinsic antimetastatic properties, further restricting their therapeutic efficacy.

Mesoporous silica nanoparticles (MSNs) provide a versatile alternative due to their tunable size, high surface area, biocompatibility, and efficient drug-loading capacity.[22] Functional modifications of MSNs allow the attachment of functional groups, enabling tailored drug loading and controlled release.[23, 24] Notably, the concept of “silicasomes”, introduced by Meng and Nel’s group, involves hybrid nanocarriers composed of lipid bilayer–coated MSNs. This design enhances biocompatibility and optimizes drug release kinetics, with silicasomes demonstrating improved pharmacokinetics, enhanced SN-38 stability, and increased tumor accumulation in multiple preclinical cancer models.[14, 25, 26] However, despite these advantages, their relatively large particle size (100–130 nm) may limit penetration into poorly vascularized or desmoplastic tumor regions, and the added complexity of lipid coating could hinder large-scale manufacturing and clinical translation.[21] To address these limitations, recent strategies have shifted toward “simpler” MSN-based systems with optimized sizes and streamlined surface modifications. For instance, MSNs can be designed as small as 25 nm, enabling deep penetration into tumor regions while maintaining high drug-loading capacity.[27] This feature facilitates access to the central hypoxic tumor areas, which is particularly critical for camptothecin-class agents such as IRI that require prolonged engagement with topoisomerase I.[6]

Our recent work further demonstrated that PEGylated MSNs functionalized with quaternary amine groups (MSN-PEG/TA) not only enhance circulation and tumor accumulation but also exhibit intrinsic antimetastatic activity by modulating focal adhesion kinase (FAK) signaling, thereby suppressing cancer cell migration and invasion.[28] This unique property suggests that MSN-based systems may offer therapeutic benefits beyond conventional drug delivery where the nanocarrier does not show any antimetastasis effect(such as liposome).

In this study, we aimed to overcome the intrinsic limitations of free IRI by developing sulfonate-functionalized mesoporous silica nanoparticles ((SO_3_^-^)-MSN-PEG/TA) as a unique nanoplatform. Under acidic conditions, these nanoparticles acquire a negatively charged surface that facilitates the loading of positively charged IRI while preserving its active lactone form. We hypothesized that this nanoplatform could simultaneously enhance IRI stability and delivery for effective suppression of primary tumor growth, while also leveraging the intrinsic antimetastatic activity of MSNs to block metastatic dissemination. Encapsulation of IRI into (SO_3_^-^)-MSN-PEG/TA (IRI@(SO_3_^-^)-MSN-PEG/TA) significantly inhibited both primary tumor growth and metastatic spread in orthotopic HCT-116 colorectal tumor models, which are characterized by highly metastatic tumors in adjacent tissues and distant organs. Both in vitro and in vivo studies demonstrated that IRI@(SO_3_^-^)-MSN-PEG/TA enhances apoptosis, suppresses tumor progression, reduces systemic toxicity, and improves overall survival by combining the cytotoxic effects of IRI with the intrinsic antimetastatic activity of MSNs, outperforming free IRI and Onivyde. IRI@(SO_3_^-^)-MSN-PEG/TA favorablyaltered the pharmacokinetic profile of IRI, prolonging systemic retention and maintaining higher IRI levels compared with free IRI. This dual-functional platform addresses critical unmet needs in mCRC therapy by optimizing key parameters: (1) enhanced tumor targeting via the EPR effect, (2) preservation of the active lactone form, (3) inhibition of metastatic dissemination, and (4) reduction of systemic toxicity through controlled drug encapsulation and release.

## 2. Materials and Methods

### 2.1. Materials

Rhodamine B isothiocyanate (RITC), (3-Mercaptopropyl)trimethoxysilane (95%), and hydrogen peroxide solution (35% (w/w) in H_2_O) were purchased from Sigma-Aldrich (Milwaukee, WI). Cetyltrimethylammonium bromide (CTAB, 99+%), tetraethyl orthosilicate (98%), ammonium hydroxide (NH_4_OH, 28-30 wt% as NH_3_), and hydrochloric acid (for analysis, fuming, 37% solution in water) were obtained from ACROS. 2-[Methoxy(polyethyleneoxy)6-9propyl]trimethoxysilane, tech-90 (PEG-silane, MW 460-590 g/mol), N-trimethoxysilylpropyl-N,N,Ntrimethylammonium chloride (TA-silane, 50% in methanol) were acquired from Gelest (Morrisville, PA). Ethanol (99.5% and 95%) was purchased from Choneye Pure Chemicals. Dulbecco’s phosphate-buffered saline (PBS) was obtained from Invitrogen. Irinotecan (IRI) was supplied by ScinoPharm Taiwan Ltd. The Cell Counting Kit-8 (CCK-8) was purchased from Abcam. The FITC Annexin V Apoptosis Detection Kit with PI was procured from BioLegend. Acetonitrile (ACN) was purchased from Honeywell and trifluoroacetic acid (TFA) were obtained from Alfa. The dialysis membrane (regenerated cellulose, MWCO 12–14 kDa) was sealed with SpectraPor® closures (Spectrum Laboratories, USA). Antibodies against phosphorylated FAK (p-FAK) and paxillin were obtained from Cell Signaling Technology, while GAPDH and secondary IgG antibodies were purchased from Santa Cruz Biotechnology. Alexa Fluor 488-conjugated secondary antibodies (green) and dimethyl sulfoxide (DMSO) were obtained from Thermo Fisher Scientific. DAPI was sourced from BioShop (Canada). TBST buffer and Triton X-100 were purchased from Sigma-Aldrich. Formalin (10%) and reagents for hematoxylin and eosin (H&E) staining were procured from Merck. Fluorescein isothiocyanate-dextran (FITC-dextran, average mol wt 70,000) was purchased from Sigma-Aldrich.

### 2.2. Synthesis of Functionalized MSNs (MSN-PEG/TA and (SO_3_^-^)-MSN-PEG/TA)

In brief, 0.29 g of CTAB was dissolved in 150 mL of 0.128 M ammonium hydroxide solution at 60°C in a sealed beaker. After 15 minutes of stirring, 0.88 M tetraethoxysilane (TEOS) dissolved in ethanol was added. After one hour of reaction, 550 μL of PEG-silane and 85.7 μL of TA-silane in 2 mL of ethanol were introduced. The mixture was stirred for an additional hour and then aged without stirring overnight. The resulting particles were subjected to hydrothermal treatment at 70°C for one day. The synthesized samples were washed and collected by centrifugation. To remove the surfactant from the MSN pores, the samples were incubated in 50 mL of acidic ethanol containing 848 μL of hydrochloric acid (37%) for the first extraction and 50 μL for the second extraction, each for one hour at 60°C. The products were washed, harvested by centrifugation, and finally stored in ethanol. The resulting particles were designated as MSN-PEG/TA.

The procedure was similar for synthesizing sulfonate-functionalized MSNs ((SO_3_^-^)-MSN-PEG/TA), with the ammonia concentration adjusted to 0.096 M at 50°C. After CTAB was dissolved for 15 minutes, a mixture of 40.6 μL of (3-mercaptopropyl)trimethoxysilane and 312 μL of TEOS in 1.4 mL ethanol (TEOS/thiol-silane ratio of 8.3:1) was added and stirred for 60 minutes. Subsequently, an additional 78 μL of TEOS in 1.2 mL ethanol was introduced. After two hours of reaction, a mixture of 1 mL of PEG-silane and 155.8 μL of TA-silane in 3.2 mL ethanol was added. The reaction mixture was stirred for one hour, aged overnight, and then subjected to hydrothermal treatment at 70°C for one day. The surfactant was removed using hydrochloric acid, and the particles were collected by centrifugation and stored in water. To convert the mercapto (thiol) groups to sulfonate groups, the synthesized MSNs (100 mg/mL) were diluted with H_2_O_2_ and stirred for over 16 hours at room temperature. The sample was then washed with H_2_O, neutralized with NaOH, washed with acetate buffer (2.5 mM, pH 5.5), and stored at 4°C.

### 2.3. Characterization of Functionalized MSNs

Transmission electron microscopy (TEM) images were obtained using a Hitachi H-7700 operating at 100 kV. Nanoparticle size distribution analysis was performed using Sigma Scan Pro 5.0 software (Ashburn, VA). Dynamic light scattering (DLS) measurements of MSN-PEG/TA and (SO_3_^-^)-MSN-PEG/TA suspended in PBS buffer were conducted using a Nano ZS90 laser particle analyzer (Malvern Instruments, U.K.). Zeta potentials of MSN-PEG/TA and (SO_3_^-^)-MSN-PEG/TA at a concentration of 0.1 mg/mL were measured in diluted PBS solution. The N₂ adsorption-desorption isotherms of MSN-PEG/TA and (SO_3_^-^)-MSN-PEG/TA were obtained using a Micrometrics ASAP 2020 system (Norcross, GA). The surface area and pore size were calculated using the Brunauer−Emmett−Teller (BET) equation and the standard Barrett−Joyner−Halenda (BJH) method. The carbon, nitrogen, sulfur, and hydrogen percentages in the dried sample (5-10 mg) were measured with an elemental analyzer (Elementar Vario EL cube type for NCSH, Germany). Thermogravimetric analysis (TGA) was recorded from 40°C to 800°C on a thermal analyzer with a heating rate of 10°C/min with an air purge of 40 mL/min.

### 2.4. Preparation of Irinotecan Loaded MSN-PEG/TA and (SO_3_^-^)-MSN-PEG/TA

250 mg of MSN-PEG/TA or (SO_3_^-^)-MSN-PEG/TA was dispersed in 2.5 mL of acetate buffer (2.5 mM, pH 4.5-5.5). A solution of 375 μL of IRI in DMSO at a concentration of 100 mg/mL was slowly added to the nanoparticle suspension with vigorous stirring for 30 minutes at room temperature. After thorough mixing, free IRI aggregates were removed by filtration through a 0.22 μm filter. The resulting solution was washed with 7- to 10-fold volumes of acetate buffer (2.5 mM, pH 4.5-5.5) to remove unbound drug and stored at 4°C.

### 2.5. IRI Quantification using High-Performance Liquid Chromatography (HPLC)

The amount of loaded IRI in IRI@MSN-PEG/TA or IRI@(SO_3_^-^)-MSN-PEG/TA was determined using high-performance liquid chromatography (HPLC). 0.2 mg sample of stock solution in 160 μL of H_2_O was mixed with 48 μL of aqueous hydrofluoric acid (HF, 1.5%) and 52 μL of acetonitrile (ACN). The mixture was sonicated for 10 minutes at room temperature and then centrifuged at 12,000 rpm for 10 minutes. 100 μL aliquot of the supernatant was subjected to HPLC analysis using a 1260 Infinity II LC System (Agilent). The HPLC conditions were as follows: a Sepax Bio-C18 column (4.6 mm × 250 mm, 5 μm particle size) was used, with a flow rate of 1.0 mL/min and a linear gradient elution from 23% solvent A to 40% solvent A. Solvent A consisted of 99.9% water with 0.1% trifluoroacetic acid (TFA), while solvent B consisted of 90% acetonitrile with 9.9% water and 0.1% TFA. The detection wavelength was set at 255 nm. The loading capacity was calculated as the weight ratio of IRI to IRI@(SO_3_^-^)-MSN-PEG/TA, and the loading efficiency was calculated as the weight ratio of the IRI loaded in IRI@(SO_3_^-^)-MSN-PEG/TA to the total amount of IRI used during the loading process.

### 2.6. Analysis of IRI Lactone Form Ratio in Free IRI and IRI@(SO_3_^-^)-MSN-PEG/TA

To evaluate the interconversion of the carboxylate and lactone forms of IRI in free IRI and IRI@(SO_3_^-^)-MSN-PEG/TA, samples were prepared in PBS at a concentration of 1 mg/mL and incubated at 37°C. The lactone form ratio [ (area from lactone form / total area from lactone and carboxylate form) x 100% ] was determined using HPLC at predetermined time points. For the analysis of free IRI, 18 µL sample was taken and mixed with 222 µL of H_2_O and 60 µL of ACN before HPLC analysis. For the analysis of IRI@(SO_3_^-^)-MSN-PEG/TA (both free and entrapped forms), 6 µL sample was mixed with 294 µL of H_2_O for HPLC analysis. For the analysis of IRI in IRI@(SO_3_^-^)-MSN-PEG/TA (entrapped form), 200 µL of the sample was gotten and washed four times with 500 µL of H_2_O using a Vivaspin concentrator. The resulting concentrate was then prepared for HPLC analysis.

### 2.7. In Vitro Drug Release

The release of IRI from IRI@(SO_3_^-^)-MSN-PEG/TA was evaluated using a dialysis method. IRI@(SO_3_^-^)-MSN-PEG/TA was suspended in 4 mL of PBS at a concentration of 1 mg IRI/mL and loaded into a dialysis membrane (regenerated cellulose, MWCO 12–14 kDa), which was sealed with SpectraPor® closures. The dialysis membrane was submerged in a 100 mL glass bottle containing 96 mL of the release medium (phosphate-buffered saline, pH 7.4) along with a stir bar. The bottle was placed in a water bath maintained at 37°C with continuous stirring.

At predetermined time intervals, 300 µL of the release medium was sampled and replaced with an equal volume of fresh release medium. To quantify the amount of IRI released into the medium, 100 µL of the collected solution was subjected to HPLC analysis.

### 2.8. Degradation Behavior of (SO_3_^-^)-MSN-PEG/TA

(SO_3_^-^)-MSN-PEG/TA was dispersed in PBS buffer solution at a concentration of 0.2 mg/mL and incubated at 37°C for several days. The morphology, hydrodynamic size, and count rate of the nanoparticles in the solution were analyzed using TEM and DLS measurements on days 0 through 7 after incubation.

### 2.9. In Vitro Cytotoxicity Assessment

HCT-116 cell lines were cultured in RPMI 1640 medium supplemented with 10% fetal bovine serum, 100 U/mL penicillin, and 100 μg/mL streptomycin, and maintained in a humidified atmosphere of 5% CO₂ at 37°C. For the cell viability assay, cells were seeded at a density of 5.0 × 10³ cells per well in a 96-well plate and incubated for 24 hours. Subsequently, the cells were treated with IRI at concentrations of 0, 6.25, 12.5, 25, 50, 100, and 200 μg/mL, IRI@(SO_3_^-^)-MSN-PEG/TA (equivalent to the various concentrations of IRI), and (SO_3_^-^)-MSN-PEG/TA (equivalent to the concentration of IRI@(SO_3_^-^)-MSN-PEG/TA). After 24 hours of incubation, cell viability was assessed using the Cell Counting Kit-8 (CCK-8). Control cells were incubated in the culture medium without treatment. The absorbance of the blank solution (100 μL of CCK-8 reagent in medium) was subtracted from the absorbance values of both the control and treated samples. All experiments were performed in triplicate, and cell viability was calculated using the formula:

Cell viability (%) = [(A sample - A Blank) / (A control - A Blank)] × 100%

### 2.10. Hemolysis Analysis

Healthy BALB/c mice (BioLasco, Taiwan), seven weeks old, were used to isolate red blood cells (RBCs) from whole blood samples. The diluted RBC suspension was mixed with (SO_3_^-^)-MSN-PEG/TA or IRI@(SO_3_^-^)-MSN-PEG/TA solutions at various concentrations (ranging from 12.5 to 1600 μg/mL). Positive control samples were incubated with water, while negative control samples were incubated with PBS solution. All samples were incubated at room temperature in the dark for 24 hours. Following incubation, the samples were centrifuged, and the absorbance of the supernatant was measured at 570 nm. The percentage of RBC hemolysis was calculated using the formula:

Hemolysis (%) = (absorbance sample - absorbance negative)/( absorbance positive - absorbance negative) x 100

### 2.11. In Vitro Cellular Uptake

HCT-116 cells were seeded in a 6-well plate at a density of 8 × 10⁵ cells per well and treated with different concentrations (250, 500, 750, and 1000 μg/mL) of R-(SO_3_^-^)-MSN-PEG/TA for 24 hours. The cellular uptake of R-(SO_3_^-^)-MSN-PEG/TA was analyzed using flow cytometry.

### 2.12. Apoptosis Detection

Cell apoptosis was quantified using the Annexin V-FITC Apoptosis Detection Kit according to the manufacturer’s instructions. Briefly, HCT-116 cells were seeded at a density of 1.5 × 10⁴ cells per well in a 24-well plate and treated with IRI (37.8 μg/mL, IC50 value), IRI@(SO_3_^-^)-MSN-PEG/TA (equivalent to 37.8 μg/mL of IRI), and (SO_3_^-^)-MSN-PEG/TA (equivalent to the concentration of IRI@(SO_3_^-^)-MSN-PEG/TA) for 24 hours. The cells were harvested with the total culture medium, washed twice with cold PBS by centrifugation, and resuspended in 100 μL of Annexin V binding buffer. The cells were then stained with 5 μL of Annexin V-FITC and 10 μL of Propidium Iodide (PI) solution in the dark for 15 minutes at room temperature, followed by the addition of 400 μL of Annexin V binding buffer. Flow cytometry was used to analyze the early apoptotic (Annexin V+/PI−) and late apoptotic (Annexin V+/PI+) cells.

### 2.13 Western Blot Analysis

HCT-116 cells were treated with (SO_3_^-^)-MSN-PEG/TA (200 μg/mL) for 24 hours and subsequently lysed in RIPA buffer containing protease and phosphatase inhibitors. Protein concentrations were quantified using a BCA assay. Equal amounts of protein (20 μg) were denatured, separated on a 10% SDS-PAGE gel, and transferred onto a PVDF membrane. The membrane was blocked with 5% BSA in TBST, followed by overnight incubation at 4°C with primary antibodies (p-FAK from Cell Signaling Technology and GAPDH from Santa Cruz Biotechnology). The membrane was then incubated with a secondary IgG antibody (Santa Cruz Biotechnology) at room temperature for 1 hour. Protein bands were visualized with an enhanced chemiluminescence substrate kit (Amersham Pharmacia Biotech, GE Healthcare UK, Bucks, UK) according to the manufacturer’s protocol.

### 2.14. Immunofluorescence Assay and Quantification

HCT-116 cells were treated with (SO_3_^-^)-MSN-PEG/TA (200 μg/mL) for 24 hours, washed twice with PBS, fixed in 4% PFA for 10 minutes, and permeabilized with 0.1% Triton X-100 for 10 minutes. After blocking with 3% BSA for 1 hour, cells were incubated overnight at 4°C with primary antibodies against paxillin (Cell Signaling Technology). Following four washes with TBST, cells were stained with Alexa Fluor 488-conjugated secondary antibodies (green) for 1 hour at room temperature. Nuclei were counterstained with DAPI (BioShop, Canada). Fluorescence images were captured using a Stellaris 8 microscope (Leica), and the number of focal adhesions per cell was quantified (n = 5).

### 2.15. Anti-Angiogenesis in the Chick Chorioallantoic Membrane (CAM)

The CAM assay was conducted to evaluate the in vivo anti-angiogenic activity of (SO_3_^-^)-MSN-PEG/TA. Fertilized eggs were incubated at 37 °C with 60% humidity for 10 days. A small window (∼1 cm in diameter) was carefully created on the eggshell using a Dremel 3000 tool. A Teflon O-ring was positioned over the CAM membrane artery, and HCT-116 cells (2.5 × 10⁶ cells in 20 μL PBS) were seeded within the ring. The window was then sealed with Tegaderm film (3M, USA), and the eggs were returned to the incubator to promote solid tumor growth. Two days after implantation (embryonic day 12), the Teflon O-ring was removed. On embryonic day 13, (SO_3_^-^)-MSN-PEG/TA (1 mg/egg) was administered intravenously into the CAM vein through the broadside opening using a syringe. Images of the vascular plexus on the CAM were captured with an optical microscope on embryonic day 15. Blood vessel density was quantified using NIH ImageJ software with the “angiogenesis analyzer” plug-in (n = 5).

### 2.16. Tumor-Targeting Dynamics of (SO_3_⁻)-MSN-PEG/TA in a Heterotopic HCT-116 Xenograft Mouse Model

Human colorectal cancer HCT-116 cells (5 × 10⁶) were subcutaneously implanted into the left flank of NOD-SCID mice to establish a heterotopic xenograft model. Mice were intravenously injected with RITC-conjugated (SO_3_⁻)-MSN-PEG/TA (200 mg/kg), and tumor regions were imaged 24 hours post-injection using multiphoton laser scanning microscopy (LSM; Olympus FVMPE-RS) equipped with a tunable infrared laser (700–1080 nm). To visualize blood vessels, 60 μL of 2.5% (w/v) fluorescein isothiocyanate–dextran (FITC–dextran, Mₙ ≈ 70 kDa) in sterile saline was administered intravenously via the tail vein. After 30 min, the mice were euthanized, and images of the tumor regions were acquired.

### 2.17. Evaluation of Dose-Dependent Efficacy of IRI@(SO_3_^-^)-MSN-PEG/TA or IRI in a Heterotopic HCT-116 Xenograft Mouse Model

HCT-116 were subcutaneously implanted into NOD-SCID mice to establish heterotopic xenografts. Once tumors reached over 200 mm³ (on Day 19), mice were received IRI or IRI@(SO_3_^-^)-MSN-PEG/TA at dose levels of 20 mg IRI/kg and 40 mg IRI/kg, administered intravenously twice per week for six total administrations. Tumor size and body weight were monitored throughout the study period (n = 5).

### 2.18. Antitumor and Antimetastasis Activity of IRI@(SO_3_^-^)-MSN-PEG/TA or (SO_3_^-^)-MSN-PEG/TA in Orthotopic Metastatic HCT116 Colorectal Tumor Model

Luciferase-expressing HCT-116 human colorectal cancer cells (2 × 10^6^) were injected into the cecum wall of NOD-SCID mice to establish an orthotopic metastatic colorectal cancer model. This model simulated the progression of highly metastatic tumors in adjacent intestinal tissues and distant organs, including the spleen, kidney, and liver.[29] Tumor-bearing mice were treated intravenously with IRI (40 mg IRI/kg), IRI@(SO_3_^-^)-MSN-PEG/TA (equivalent to 40 mg IRI/kg), and (SO_3_^-^)-MSN-PEG/TA (equivalent to the concentration of IRI@(SO_3_^-^)-MSN-PEG/TA). Treatments began on Day 7 and were administered intravenously twice a week for eight administrations. Tumor growth and metastatic progression were monitored weekly using an in vivo imaging system (IVIS) to capture bioluminescence signals from the luciferase-expressing cells. Body weight was recorded throughout the study (n = 5). Mice were closely observed for overall survival, with monitoring until spontaneous death or the onset of a moribund state.

### 2.19. In Vivo Toxicity and Histological Analysis

Male BALB/c mice, 7 weeks old (N=4), were administered IRI alone (40 and 60 mg IRI/kg), IRI@(SO_3_^-^)-MSN-PEG/TA (equivalent to the concentration of IRI), (SO_3_^-^)-MSN-PEG/TA (equivalent to the concentration of IRI@(SO_3_^-^)-MSN-PEG/TA and Onivyde (equivalent to the concentration of IRI) via intravenous tail vein injection every day for 4 administrations. Body weight was recorded every 4 days. After the final administration, the mice were sacrificed, and serum was isolated from whole blood for complete blood count (CBC) and blood chemistry (BC) analyses, which were conducted by the Taipei Medical University Laboratory Animal Center. To study the toxicological pathology, organs such as the liver, sternum, and intestine were fixed with 10% formalin, embedded in paraffin, and sectioned. Hematoxylin and eosin (H&E) staining were performed on the tissue sections for histological analysis. An experienced veterinary pathologist (TOSON Technology Co., Ltd.) evaluated the tissue slides.

### 2.20. Pharmacokinetic Study

Healthy Male Wistar rats (7 weeks old) received a single intravenous injection of either free IRI or IRI@(SO_3_^-^)-MSN-PEG/TA at a dose of 40 mg IRI/kg. Blood samples (≈100 μL) were collected via the tail vein at 0.17, 1, 2, 4, 8, and 24 hours’ post-injection into K₂EDTA anticoagulant tubes. Plasma was separated by centrifugation at 3,000 rpm for 10 min at 4°C and stored at –80°C until analysis. Camptothecin (CPT) was added as an internal standard before extraction. Plasma proteins were precipitated with an eightfold volume of extraction solvent (50% acetonitrile / 50% methanol, v/v), followed by sonication for 15 min and centrifugation at 14,000 rpm for 15 min. The supernatant was collected and diluted to a final composition of 30% organic solvent (15% acetonitrile, 15% methanol, and 70% water), then centrifuged again at 14,000 rpm for 15 min prior to analysis.

Quantification was performed using an Agilent 6470 Triple Quadrupole mass spectrometer equipped with an electrospray ionization (ESI) source, coupled to an Agilent 1260 Infinity II Quaternary Pump LC system. Chromatographic separation was achieved on an ACQUITY UPLC BEH C18 column (2.1 mm × 100 mm, 1.7 μm, 130 Å; Waters, SKU#186002350) using water (solvent A) and acetonitrile (solvent B), each containing 0.1% formic acid, as the mobile phase at a flow rate of 0.4 mL/min. The gradient was programmed as follows: 5% B for 1 min, ramped to 100% B over 8 min, held for 1 min, then returned to 5% B at 8.1 min, with a total run time of 10 min. The column temperature was maintained at 40 °C, and the injection volume was 2 μL with the autosampler at room temperature.

The mass spectrometer was operated in multiple reaction monitoring (MRM) acquisition mode under positive electrospray ionization (ESI⁺). The protonated ions of IRI [M+H]⁺, and CPT [M+H]⁺ were monitored at m/z 587.3 → 167.1, and 349.1 → 305.1, respectively. Pharmacokinetic parameter values, including C_max_, AUC_₀–∞_, t_1/2_, clearance (CL), and volume of distribution (Vd), were based on the plasma concentrations of total irinotecan (including free + MSN-loaded irinotecan) and determined using the non-compartmental analysis with PKSolver, an Excel-based pharmacokinetic add-in.

## 3. Results and Discussion

### 3.1. Customized Design of Mesoporous Silica Nanoparticles for Enhanced Irinotecan Loading

Developing efficient and targeted drug delivery systems is essential for advancing mCRC therapy. Building on our previous findings that PEGylated MSNs functionalized with quaternary amine groups (MSN-PEG/TA) possess intrinsic antimetastatic and tumor-targeting properties[28], we further optimized these nanoparticles for IRI delivery. To enhance electrostatic interactions with protonated IRI under acidic conditions, sulfonate groups were introduced into the inner pores, generating(SO_3_^-^)-MSN-PEG/TA.

Transmission electron microscopy (TEM) revealed that MSN-PEG/TA and (SO_3_^-^)-MSN-PEG/TA exhibited average diameters of 28.5 ± 4.5 nm and 25.5 ± 4.0 nm, respectively (Figure 1a-c and Table S1). Both nanoparticles showed hydrodynamic diameters of approximately 40 nm in PBS, with polydispersity indices (PDI) below 0.1, indicating excellent dispersion (Figure 1d). Zeta potential measurements confirmed the effects of surface modifications (Figure 1e and Table S1). MSN-PEG typically exhibited a negative surface charge (≈ –15 mV, data not shown), attributable to silica surface silanol groups. After modification with positively charged TA-silane, MSN-PEG/TA displayed a less negative potential (–8.1 ± 0.5 mV), reflecting partial charge neutralization by quaternary amine groups. Incorporation of sulfonate groups significantly shifted the potential to −28.2 ± 0.8 mV, corroborated by elemental analysis showing 2.25 ± 0.1% sulfur content in (SO_3_^-^)-MSN-PEG/TA, with no detectable sulfur in MSN-PEG/TA (Table S2). Nitrogen adsorption–desorption isotherms exhibited typical type IV behavior with a pronounced hysteresis loop, which is characteristic of mesoporous materials (Figure 1f). The Brunauer–Emmett–Teller (BET) surface areas were 325.9 m²/g for MSN-PEG/TA and 311.9 m²/g for (SO_3_^-^)-MSN-PEG/TA, with average pore diameters of 1.5 and 1.36 nm, respectively, calculated by the Barrett–Joyner–Halenda (BJH) method. Thermogravimetric analysis (TGA) validated successful surface modification (Figure 1g and Table S3). Three distinct stages were observed between 40°C and 800°C: (1) 40–200°C, loss of physically adsorbed water; (2) 200–600°C, decomposition of surface-grafted organic groups (PEG, TA, and sulfonic acid); and (3) 600–800°C, degradation and carbonization of residual organics. Weight loss in the 200–600°C region was 24.9% for MSN-PEG, 33.9% for MSN-PEG/TA, and 37.5% for (SO_3_^-^)-MSN-PEG/TA, compared to only 2.4% for bare MSN. The progressive increase in weight loss reflects the higher content of surface-bound organic moieties, with an additional 3.6% in (SO_3_^-^)-MSN-PEG/TA confirming successful sulfonate incorporation.

**Figure 1.**
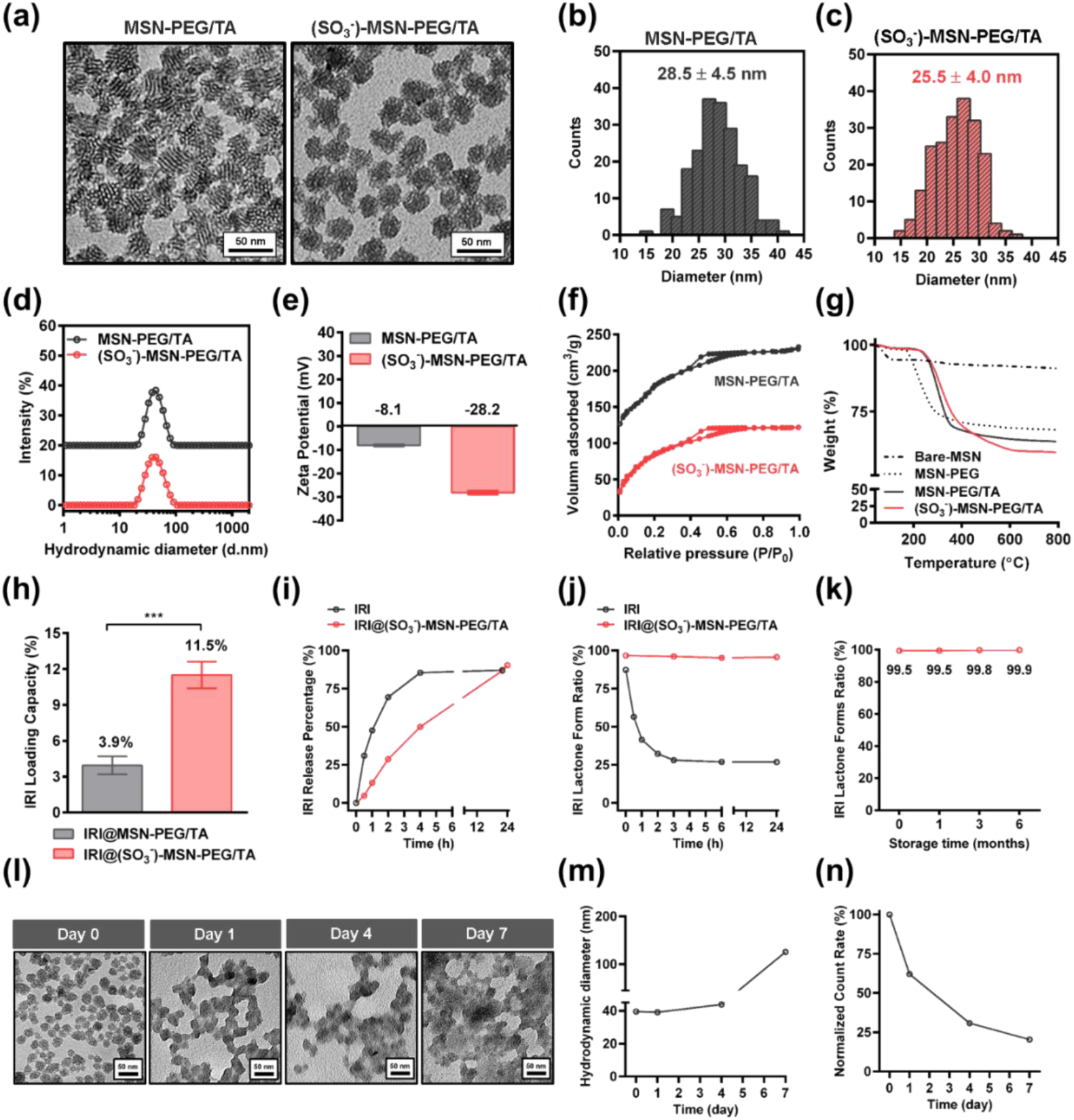
Characterization of functionalized MSNs. (a) TEM images of MSN-PEG/TA and (SO_3_^-^)-MSN-PEG/TA. Scale bar = 50 nm. (b-c) The panels show particle size distributions obtained from TEM image analysis. (d) Hydrodynamic diameters and (e) zeta potentials of MSN-PEG/TA and(SO_3_^-^)-MSN-PEG/TA measured by DLS. (f) N₂ adsorption–desorption isotherms of MSN-PEG/TA and (SO_3_^-^)-MSN-PEG/TA showing BET surface areas and pore sizes (BJH method). (g) Thermogravimetric analyses of bare MSN, MSN-PEG, MSN-PEG/TA, and (SO_3_⁻)-MSN-PEG/TA. Detailed results are summarized in Table S3. (h) Comparison of IRI-loading performance between MSN-PEG/TA and (SO_3_^-^)-MSN-PEG/TA, expressed as loading capacity (LC%). ***p < 0.001, n = 5. (i) In vitro drug release profiles of free IRI and IRI@(SO_3_^-^)-MSN-PEG/TA. (j) Time-dependent ratio of IRI’s active lactone form to the carboxylate form over time for free IRI and IRI@(SO_3_^-^)-MSN-PEG/TA in PBS. (k) Stability of the lactone/total IRI ratio in IRI@(SO_3_^-^)-MSN-PEG/TA after six months of storage in acetate buffer (2.5 mM, pH 5.5). (l) In vitro degradation of (SO_3_⁻)-MSN-PEG/TA in PBS at 37 °C for 7 days, with TEM images showing progressive morphological changes. (m) DLS analysis of (SO_3_^-^)-MSN-PEG/TA at different time points demonstrating size and (n) count rate variations over time. Scale bar = 50 nm.

A key distinction between the two MSNs was observed in their IRI loading capacity and efficiency under acidic conditions (pH ≤ 5.5). Due to its weakly negative surface charge, MSN-PEG/TA exhibited limited loading (capacity 3.9%, efficiency 45%). In contrast, (SO_3_^-^)-MSN-PEG/TA achieved a markedly higher loading capacity of 11.5% and efficiency exceeding 90% (Figure 1h and Table S1). IRI becomes protonated and positively charged in acidic environments, while sulfonic groups (SO₃⁻, pKa ≈ –7) remain strongly negative charge. This electrostatic interaction enables efficient drug loading and preserves IRI in its active lactone form within the acidic tumor microenvironment. These results identify (SO_3_^-^)-MSN-PEG/TA as a superior carrier for IRI encapsulation. The IRI release profile was evaluated in PBS at 37 °C. Free IRI displayed rapid release, reaching 80% within 4 hours, whereas IRI@(SO_3_^-^)-MSN-PEG/TA exhibited sustained release over 20 hours (Figure 1i). Conversion between the active lactone and inactive carboxylate forms of IRI was monitored by high-performance liquid chromatography (HPLC). Free IRI rapidly converted to the inactive carboxylate form within 3 hours, while IRI@(SO_3_^-^)-MSN-PEG/TA retained over 90% of the active lactone form after 24 hours (Figure 1j). Long-term storage stability was further confirmed, with IRI@(SO_3_^-^)-MSN-PEG/TA maintaining nearly 99% of the active lactone form after 6 months in acetate buffer (Figure 1k). These results demonstrate that (SO_3_^-^)-MSN-PEG/TA effectively stabilizes and preserves IRI’s active lactone form, a critical factor for maintaining therapeutic potency.

The hydrolytic degradation of (SO_3_^-^)-MSN-PEG/TA was examined by incubation in PBS at 37°C for 7 days (Figure 1l-n). TEM and DLS analyses revealed progressive structural disintegration and aggregation by day 4, with near-complete degradation by day 7. This gradual breakdown supports sustained drug release and highlights the biodegradable nature of (SO_3_^-^)-MSN-PEG/TA, advantageous for minimizing long-term systemic accumulation. In summary, the rationally engineered IRI@(SO_3_^-^)-MSN-PEG/TA combines high drug-loading efficiency, exceptional lactone-form stability, controlled release, and intrinsic biodegradability, making it a promising nanoplatform for subsequent in vitro and in vivo therapeutic evaluation.

### 3.2. Cellular Uptake, Hemolytic Activity, and Anti-Proliferative Effects of (SO_3_^-^)-MSN-PEG/TA and IRI@(SO_3_^-^)-MSN-PEG/TA

The intracellular internalization of (SO_3_^-^)-MSN-PEG/TA was assessed using rhodamine isothiocyanate (RITC)-conjugated (SO_3_^-^)-MSN-PEG/TA (R-(SO_3_^-^)-MSN-PEG/TA) in human colorectal cancer HCT-116 cells by flow cytometry. As shown in Figure S1, cellular uptake increased in a concentration-dependent manner, confirming efficient nanoparticle internalization. To assess the cytotoxicity, cell viability was examined using the CCK-8 assay after 24 h of incubation with free IRI, IRI@(SO_3_^-^)-MSN-PEG/TA (equivalent IRI doses of 0–200 μg/mL), and blank (SO_3_^-^)-MSN-PEG/TA (equivalent to IRI@(SO_3_^-^)-MSN-PEG/TA dose). Both free IRI and IRI@(SO_3_^-^)-MSN-PEG/TA induced a dose-dependent decrease in cell viability (Figure 2a). Notably, IRI@(SO_3_^-^)-MSN-PEG/TA exhibited a lower IC₅₀ value (28.2 μg/mL) compared with free IRI (37.8 μg/mL), indicating improved drug delivery efficiency mediated by (SO_3_^-^)-MSN-PEG/TA. In contrast, (SO_3_^-^)-MSN-PEG/TA alone showed no cytotoxicity across all tested concentrations, confirming its high biocompatibility.

**Figure 2.**
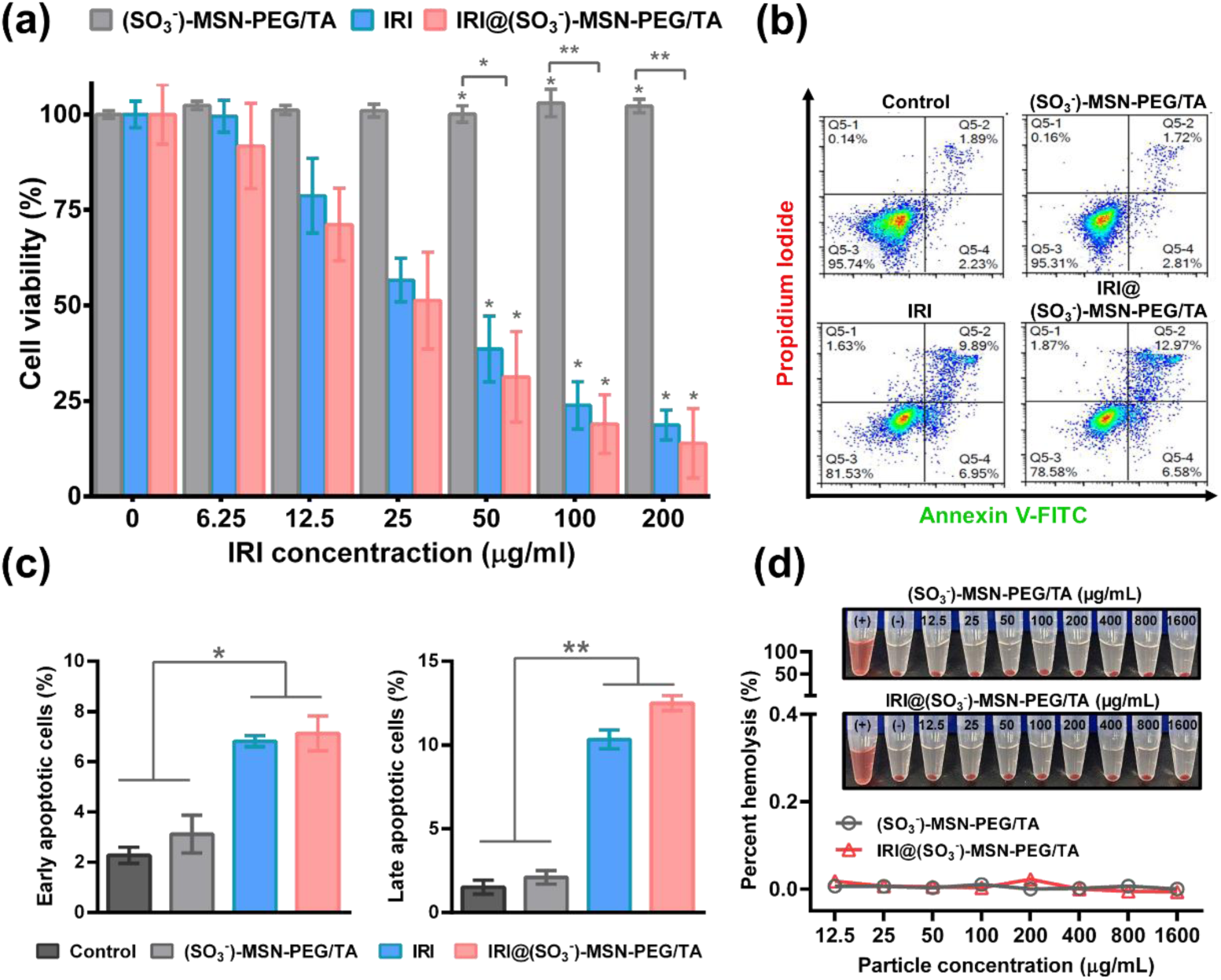
Evaluation of cytotoxicity, hemolysis, and apoptosis. (a) Cell viability of HCT-116 cells treated with different concentrations of IRI, IRI@(SO_3_^-^)-MSN-PEG/TA (equivalent IRI dose), and blank (SO_3_^-^)-MSN-PEG/TA (equivalent to the concentration of IRI@(SO_3_^-^)-MSN-PEG/TA) for 24 hours. *p < 0.05, **p < 0.01, n = 5. (b-c) Apoptosis analysis of HCT-116 cells treated with IRI (37.8 μg/mL), IRI@(SO_3_^-^)-MSN-PEG/TA (equivalent IRI dose), and (SO_3_^-^)-MSN-PEG/TA (equivalent to the concentration of IRI@(SO_3_^-^)-MSN-PEG/TA) for 24 hours, using the Annexin V/PI apoptosis detection kit. *p < 0.05, **p < 0.01, n = 5. (d) Hemolysis assay of red blood cells (RBCs) incubated with varying concentrations (1–1600 μg/mL) of (SO_3_^-^)-MSN-PEG/TA or IRI@(SO_3_^-^)-MSN-PEG/TA for 24 hours. Hemolysis was quantified by measuring hemoglobin release from lysed RBCs in the supernatant. Water (+) and PBS (−) served as positive and negative controls, respectively. n = 3.

IRI-induced apoptosis was further evaluated using an Annexin V/PI staining followed by quantitative flow cytometry analysis (Figure 2b-c). Both IRI and IRI@(SO_3_^-^)-MSN-PEG/TA induced significant apoptosis compared with the control group. The proportion of early apoptotic cells (Annexin V⁺/PI⁻) significantly increased with IRI of 6.94 ± 0.22% and IRI @(SO_3_^-^)-MSN-PEG/TA of 7.91 ± 0.7%, supporting a consistent pro-apoptotic effect. Importantly, IRI@(SO_3_^-^)-MSN-PEG/TA triggered a higher proportion of late apoptotic cells (Annexin V⁺/PI⁺, 12.5 ± 0.45%) than free IRI (10.34 ± 0.56%), indicating enhanced intracellular release and cytotoxic action of IRI following nanoparticle uptake.

To further confirm biosafety, particularly the effect of IRI loaded MSN formulation on blood compatibility, the hemocompatibility of both (SO_3_^-^)-MSN-PEG/TA and IRI@(SO_3_^-^)-MSN-PEG/TA was evaluated using a red blood cell (RBC) hemolysis assay. RBC suspensions were incubated with increasing nanoparticle concentrations (1–1600 μg/mL), and hemoglobin release due to RBC membrane disruption was quantified spectrophotometrically (Figure 2d). Across all concentrations, both formulations exhibited negligible hemolysis, while the water-treated positive control showed complete lysis. These results demonstrate that (SO_3_^-^)-MSN-PEG/TA does not compromise RBC membrane integrity and is therefore suitable for intravenous administration. Collectively, these findings confirm that (SO_3_^-^)-MSN-PEG/TA exhibits excellent biocompatibility and hemocompatibility, while serving as an effective and safe nanocarrier that enhances IRI delivery and pro-apoptotic efficacy in colorectal cancer cells.

### 3.3. Antimetastatic Effects and Tumor-Targeting Capability of (SO_3_^-^)-MSN-PEG/TA *In Vitro* and *In Vivo*

Our previous work demonstrated that MSN-PEG/TA could suppress metastasis by disrupting focal adhesion dynamics, a key process governing cancer cell migration and invasion.[28] Building upon these findings, we investigated whether sulfonate-functionalized nanoparticles ((SO_3_^-^)-MSN-PEG/TA) exert similar or enhanced antimetastatic activity by targeting focal adhesion–related signaling pathways. In particular, we examined two critical regulators of cell adhesion and motility: focal adhesion kinase (FAK) and paxillin.[30–36] Western blot analysis revealed a significant reduction in phosphorylated FAK (p-FAK) levels in HCT-116 cells treated with (SO_3_^-^)-MSN-PEG/TA (200 μg/mL, 24 hours) compared with untreated controls (Figure 3a). Quantitative analysis confirmed a marked decrease in p-FAK expression (Figure 3b), indicating that the nanoparticles effectively modulate this central signaling cascade. Because dynamic focal adhesion (FA) turnover relies heavily on paxillin activation, a key protein orchestrating FA assembly and disassembly,[37, 38] we next performed immunofluorescence staining to visualize paxillin. (SO_3_^-^)-MSN-PEG/TA treatment markedly reduced paxillin-positive focal adhesions (green) relative to controls, with nuclei counterstained by DAPI (blue) (Figure 3c). Quantification revealed that the number of focal adhesions per cell decreased from 35 ± 3.9 in controls to 16.2 ± 5.1 after treatment (Figure 3d). Together, these results suggest that (SO_3_^-^)-MSN-PEG/TA effectively impairs cellular adhesion and migration by modulating FAK/paxillin signaling.

**Figure 3.**
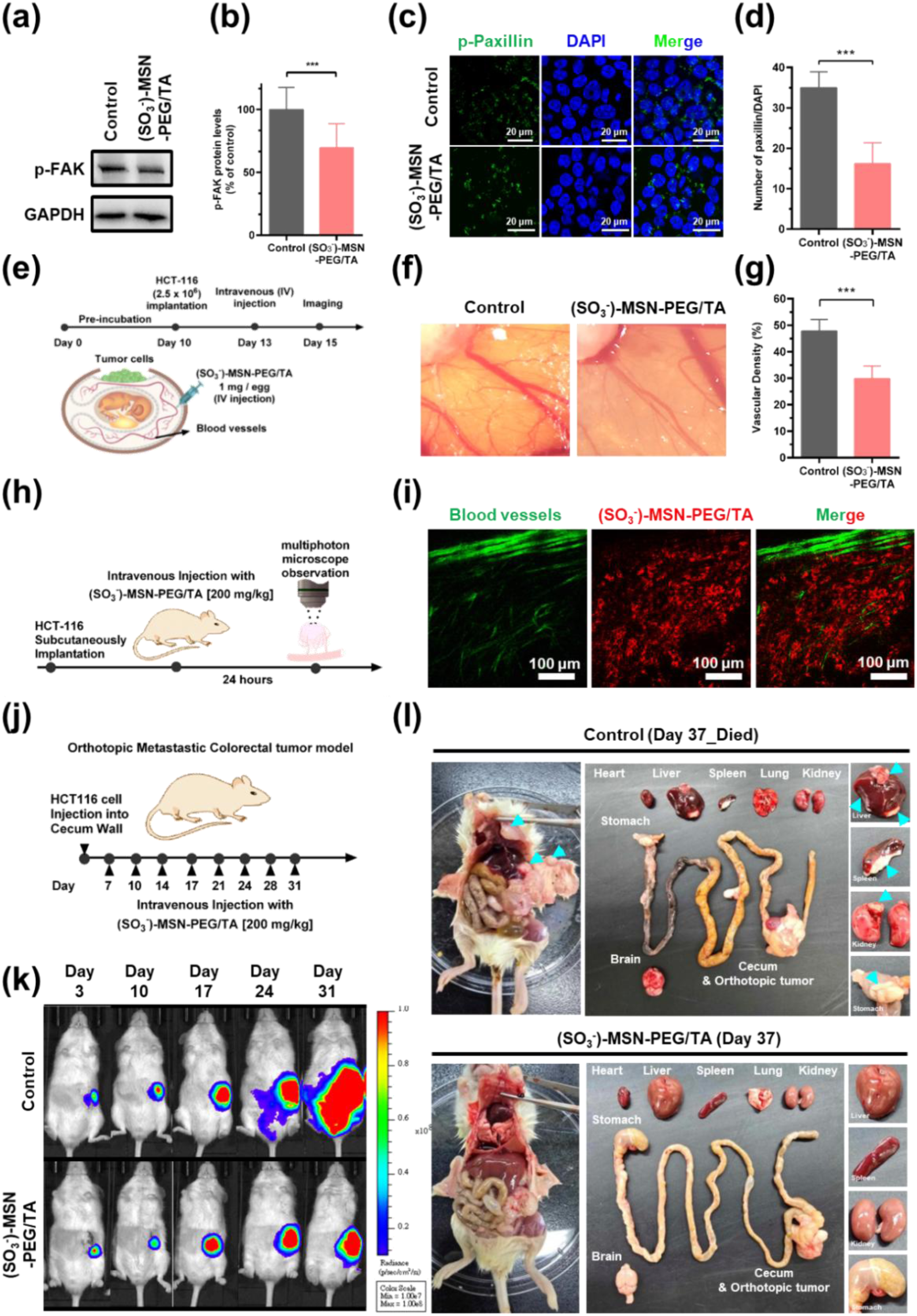
Mechanistic evaluation of (SO_3_^-^)-MSN-PEG/TA on metastasis, focal adhesion turnover, angiogenesis, and tumor targeting. HCT-116 cells were treated with 200 μg/mL of (SO_3_^-^)-MSN-PEG/TA for 24 hours. (a) Western blot analysis of p-FAK expression levels. (b) Quantitative analysis of p-FAK protein expression relative to the control group (***p < 0.001, n = 3). (c) Immunofluorescence imaging of paxillin (green) and nuclei (blue). Scale bar: 20 μm. (d) Quantitative comparison of the number of FA per cell between control and treated groups (***p < 0.001, n = 5). (e) Schematic representation of the chicken embryo CAM assay. HCT-116 cells were seeded onto the CAM, and (SO_3_^-^)-MSN-PEG/TA (1 mg/egg) was administered intravenously on embryonic day 13. (f) Representative photographs of CAM vasculature on day 15. (g) Statistical analysis of vascular density performed using ImageJ software (***p < 0.001, n =5). (h) Schematic illustration of the in vivo tumor-targeting study. HCT-116 tumor-bearing mice were intravenously injected with RITC-labeled (SO3⁻)-MSN-PEG/TA (200 mg/kg) and analyzed by multiphoton microscopy. (i) Visualization of the EPR effect. Blood vessels were labeled with FITC–dextran (green), and RITC-labeled (SO3⁻)-MSN-PEG/TA (red) signals were observed 24 hours post-injection. Scale bar = 100 μm. (j) Schematic of the orthotopic colorectal cancer metastasis model. Luciferase-expressing HCT-116 tumor tissue (2 x 10⁶ cells) was implanted into the cecum wall of NOD-SCID mice to establish an orthotopic colorectal cancer model. Mice were intravenously treated with (SO_3_^-^)-MSN-PEG/TA (equivalent to NPs dose corresponding to 40 mg/kg of IRI@(SO_3_^-^)-MSN-PEG/TA). (k) Monitoring of orthotopic tumor growth by IVIS imaging from day 3 to day 31. (l) Ex vivo imaging of metastatic lesions: Post-mortem imaging of mice revealed extensive metastasis, with tumor sites identified in organs such as the liver, spleen, kidneys, stomach, and cecum (indicated by cyan arrows).

Given the strong association between FAK activation and tumor angiogenesis, inhibition of this pathway is also expected to suppress tumor vascularization, and thereby limit tumor growth and metastasis.[30–32] To assess the anti-angiogenic activity of (SO_3_^-^)-MSN-PEG/TA in vivo, we employed the chick chorioallantoic membrane (CAM) tumor model, which enables rapid tumor formation and vascularization under physiologically relevant conditions.[39, 40] Fig. 3e illustrates the experimental setup for the CAM-based angiogenesis assay. In the control group, HCT-116 tumors (2.5 × 10⁶ cells) implanted on the CAM exhibited dense, network-like vascular growth near the tumor (Figure 3f). In contrast, embryos treated intravenously with (SO_3_^-^)-MSN-PEG/TA (1 mg/egg) displayed markedly disrupted vascular architecture and reduced blood vessel formation. Quantitative analysis of vascular density confirmed a significant decrease compared with the control group (Figure 3g; Figure S2), demonstrating potent inhibition of tumor-induced angiogenesis. These findings highlight the potential of (SO_3_^-^)-MSN-PEG/TA to function as an anti-angiogenic nanocarrier.

Efficient delivery of nanoparticle to tumor site is essential for therapeutic efficacy.[28] To evaluate tumor-targeting capability in vivo, RITC-labeled (SO_3_^-^)-MSN-PEG/TA (200 mg/kg) was intravenously administered to HCT-116 xenograft-bearing mice, and tumor regions were visualized 24 hours post-injection using multiphoton laser scanning microscopy (Figure 3h). Blood vessels were visualized using FITC–dextran (70 kDa) to assess nanoparticle extravasation. Strong red fluorescence signals corresponding to (SO3⁻)-MSN-PEG/TA were observed both adjacent to and beyond vascular boundaries, confirming extravasation and tumor accumulation through the enhanced permeability and retention (EPR) effect (Figure 3i). These results verify that (SO_3_^-^)-MSN-PEG/TA efficiently accumulates within tumor tissues, supporting its role as an effective nanocarrier for targeted therapy. Finally, the therapeutic relevance of the intrinsic antimetastatic activity of (SO_3_^-^)-MSN-PEG/TA was examined in an orthotopic colorectal cancer model. Luciferase-expressing HCT-116 tumor tissue (2×10^6^ cells) was implanted into the cecum wall of NOD-SCID mice to establish highly aggressive and metastatic colorectal tumors (Figure 3j).[14, 41–43] Mice received intravenous administration of (SO_3_^-^)-MSN-PEG/TA (equivalent to the nanoparticle dose used in IRI@(SO_3_^-^)-MSN-PEG/TA; 40 mg/kg) at scheduled intervals, and tumor progression was monitored by IVIS imaging. Compared with controls, the (SO_3_^-^)-MSN-PEG/TA treatment did not inhibit primary tumor growth but significantly suppressed cancer metastasis from day 3 to day 31 (Figure 3k). Control mice succumbed by day 37 due to extensive metastatic dissemination. Ex vivo imaging further revealed widespread metastases in the control group, including the liver, spleen, kidneys, cecum, and stomach, whereas the (SO_3_^-^)-MSN-PEG/TA-treated mice displayed tumors restricted to the primary cecal site with no detectable secondary lesions (Figure 3l, cyan arrows). Notably, the control group’s stomach was entirely covered with tumor tissue, and fat deposits in the intestines had hardened into tumor-associated nodules. Collectively, these results demonstrate that (SO_3_^-^)-MSN-PEG/TA exerts potent antimetastatic and anti-angiogenic effects by disrupting FAK/paxillin signaling and inhibiting vascular remodeling, while simultaneously achieving efficient tumor accumulation through the EPR effect. This intrinsic bioactivity positions (SO_3_^-^)-MSN-PEG/TA as an effective multifunctional nanocarrier for metastasis suppression and targeted drug delivery in colorectal cancer.

### 3.4. Inhibition of Primary Tumor Growth and Metastasis by IRI@(SO_3_^-^)-MSN-PEG/TA in a Colorectal Cancer Model

To investigate the therapeutic efficacy of IRI@(SO_3_^-^)-MSN-PEG/TA in vivo, a subcutaneous xenograft model was established by implanting HCT-116 human colorectal cancer cells (5×10^6^ cells) into NOD-SCID mice (Figure 4a). The xenograft mice received intravenous injections of free IRI or IRI@(SO_3_^-^)-MSN-PEG/TA at doses of 20 mg and 40 mg IRI/kg, administered twice weekly for six treatments. Tumor volumes and weights were recorded throughout the study. Both IRI and IRI@(SO_3_^-^)-MSN-PEG/TA treatments produced dose-dependent inhibition of tumor growth (Figure 4b). Notably, complete tumor suppression was achieved in the 40 mg IRI/kg IRI@(SO_3_^-^)-MSN-PEG/TA-treated group, whereas the same dose of free IRI only partially inhibited tumor progression. Moreover, the 20 mg/kg dose of IRI@(SO_3_^-^)-MSN-PEG/TA achieved antitumor efficacy comparable to that of 40 mg/kg free IRI, demonstrating the enhanced therapeutic potency conferred by the nanoformulation. No significant loss in body weight was observed in any treatment group (Figure 4c), confirming that IRI@(SO_3_^-^)-MSN-PEG/TA exhibits excellent biocompatibility and minimal systemic toxicity.

**Figure 4.**
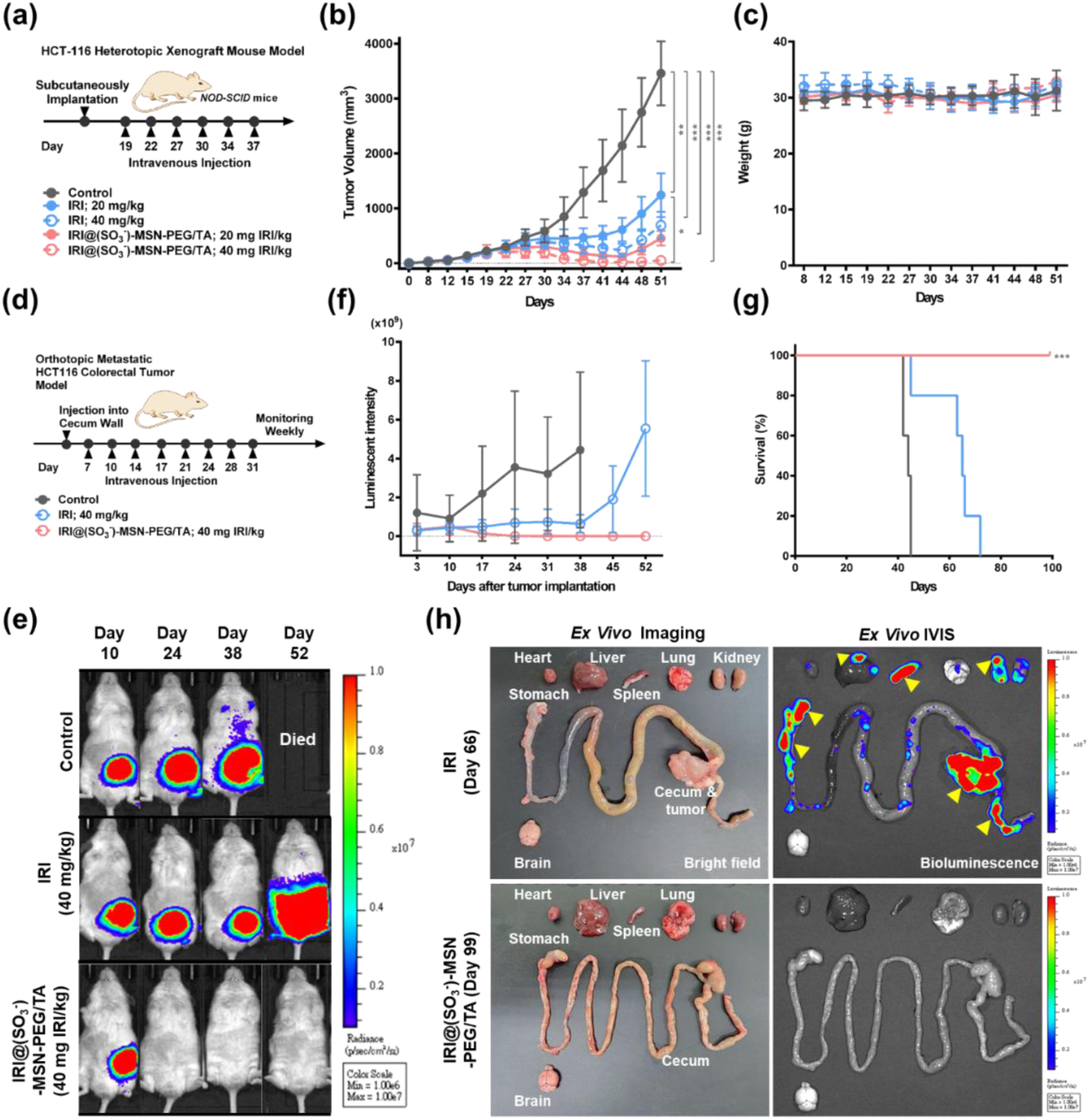
Therapeutic efficacy of IRI@(SO_3_^-^)-MSN-PEG/TA in a colorectal cancer metastasis model. (a) Schematic representation of the subcutaneous xenograft model. HCT-116 human colorectal cancer cells (5×10⁶) were implanted subcutaneously into NOD-SCID mice. Mice received intravenous injections of free IRI or IRI@(SO_3_^-^)-MSN-PEG/TA at doses of 20 mg and 40 mg IRI/kg, administered twice weekly for six treatments. (b) Tumor volume measurements and (c) body weight monitoring during the treatment period (*p < 0.05, **p < 0.01, ***p < 0.001 n = 4). (d) Schematic representation of the orthotopic colorectal cancer metastasis model, established by implanting HCT-116 tumor tissue (2×10^6^ cells) into the cecum wall of NOD-SCID mice. Mice received intravenous injections of free IRI or IRI@(SO_3_^-^)-MSN-PEG/TA (40 mg IRI/kg), administered twice weekly for a total of eight treatments. (e) Representative IVIS images showing tumor progression over time (n = 4). (f) Quantitative analysis of bioluminescence intensity from IVIS imaging, presented as mean ± SD (n = 4). (g) Kaplan-Meier survival curves comparing the survival rate among treatment groups. (***p < 0.001, n = 4) (h) Ex vivo bright-field and IVIS imaging of isolated organs from treated mice in the IRI@(SO_3_^-^)-MSN-PEG/TA group (Day 99) and the IRI group (Day 66). Yellow arrows indicate metastatic tumor sites in organs including the liver, spleen, kidney, stomach, and cecum.

Next, the therapeutic efficacy and antimetastatic potential of IRI@(SO_3_^-^)-MSN-PEG/TA were further examined using an orthotopic colorectal cancer metastasis model, in which HCT-116 tumor tissue was implanted into the cecum wall of NOD-SCID mice. The treatment schedule is outlined in Figure 4d. As shown in Figure 4e, primary tumor signals became detectable in all groups by day 10. Control mice succumbed by day 38 due to extensive metastatic dissemination. Treatment with free IRI initially suppressed primary tumor growth within the first 38 days; however, significant increase in tumor signals were observed by day 52, indicating aggressive metastatic relapse. In contrast, IRI@(SO_3_^-^)-MSN-PEG/TA treatment resulted in complete disappearance of tumor signals by day 24, consistent with substantial inhibition of primary tumor growth. Quantitative bioluminescence analysis confirmed that IRI@(SO_3_^-^)-MSN-PEG/TA effectively suppressed primary tumor progression and prevented metastasis, while free IRI failed to maintain long-term control and showed accelerated tumor regrowth after day 45 (Figure 4f). Kaplan–Meier survival analysis further revealed markedly improved survival in the IRI@(SO_3_^-^)-MSN-PEG/TA -treated group, which achieved 100 % survival, whereas all mice in the free IRI group died by approximately day 75 (Figure 4g). Although IRI is clinically approved for mCRC, it primarily functions as a cytotoxic agent that suppresses primary tumor growth but lacks intrinsic antimetastatic activity, ultimately resulting in mortality. Consistent with this limitation, control mice exhibited the shortest survival due to rapid tumor progression and widespread metastasis.

To clarify the role of metastasis in mortality, ex vivo IVIS imaging was performed on isolated organs post-mortem. As shown in Figure 4h, mice treated with IRI@(SO_3_^-^)-MSN-PEG/TA displayed a pronounced reduction in metastatic lesions compared with those treated with free IRI. Yellow arrows indicate metastatic tumor locations observed in the liver, spleen, kidney, stomach, and cecum. These results prove the excellent IRI delivery and antimetastatic capabilities of IRI@(SO_3_^-^)-MSN-PEG/TA, which may effectively concentrate IRI at the tumor site while minimizing systemic exposure. We speculate that the enhanced performance is attributed to both enhanced tumor accumulation via the EPR effect and the intrinsic antimetastatic properties of the (SO_3_^-^)-MSN-PEG/TA carrier, which collectively inhibit dissemination of cancer cells to distant organs. Overall, these findings highlight the dual therapeutic potential of IRI@(SO_3_^-^)-MSN-PEG/TA in simultaneously suppressing primary tumor growth and mitigating metastasis. This multifunctional nanoplatform provides a potential therapeutic opportunity to improve clinical outcomes in patients with mCRC.

Metastasis in cancer has many complex pathological pathways involving many targets, druggable or not. Clinical trials using metastasis-free as end point are costly. Broadly applicable anti-metastasis drugs do not yet exist. Thus we have only metastasis-directed drugs but almost no broadly applicable anti-metastasis drug. Two approved metastasis directed antibody drugs, Denosumab (against RANKL)[44] and Bevacizumab[45] are effective only to specific tumors. Unlike antibodies, MSN carrying anti-cancer drug could be a generally applicable to prevent other cancer’s metastasis while being relatively cheap. MSN’s anti-metastasis comes from binding integrin by MSN’s fragments.

Because integrin tends to be shared among large subsets of invasive cancers, a marker of enhanced invasiveness and metastatic behavior of the tumour, the development of anti-metastasis MSN-based drug may be more beneficial in this regard than the development of highly specific inhibitors.

### 3.5. In Vivo Biosafety Assessment of (SO_3_^-^)-MSN-PEG/TA and IRI@(SO_3_^-^)-MSN-PEG/TA

A comprehensive toxicity evaluation was conducted to assess the biosafety of the treatment regimens in healthy BALB/c mice, with comparisons made to the clinically approved IRI liposome (Onivyde) (Figure 5, Figures S3, and Tables S4–S5). Onivyde, approved in 2015 for the treatment of metastatic pancreatic ductal adenocarcinoma (PDAC), is known to cause severe and potentially life-threatening adverse effects such as neutropenia and diarrhea.[15, 19, 46] In this study, mice were intravenously administered IRI, IRI@(SO_3_^-^)-MSN-PEG/TA, and Onivyde at 40 and 60 mg IRI/kg, or blank (SO_3_^-^)-MSN-PEG/TA at 300 and 460 mg/kg (equivalent to the particle concentration of IRI@(SO_3_^-^)-MSN-PEG/TA), once daily for four doses (Figure 5a). As shown in Figure 5b, mice receiving the highest doses of free IRI and Onivyde (60 mg IRI/kg) exhibited significant body weight loss (>15%), indicating severe systemic toxicity. In contrast, mice treated with IRI@(SO_3_^-^)-MSN-PEG/TA (60 mg IRI/kg) displayed no abnormal behavior or weight loss, demonstrating that encapsulation of IRI in the (SO_3_^-^)-MSN-PEG/TA effectively mitigated drug-induced toxicity. Likewise, blank (SO_3_^-^)-MSN-PEG/TA caused no significant body weight changes, indicating that the nanoformulation alleviated systemic toxicity while preserving tumor-suppressive efficacy.

**Figure 5.**
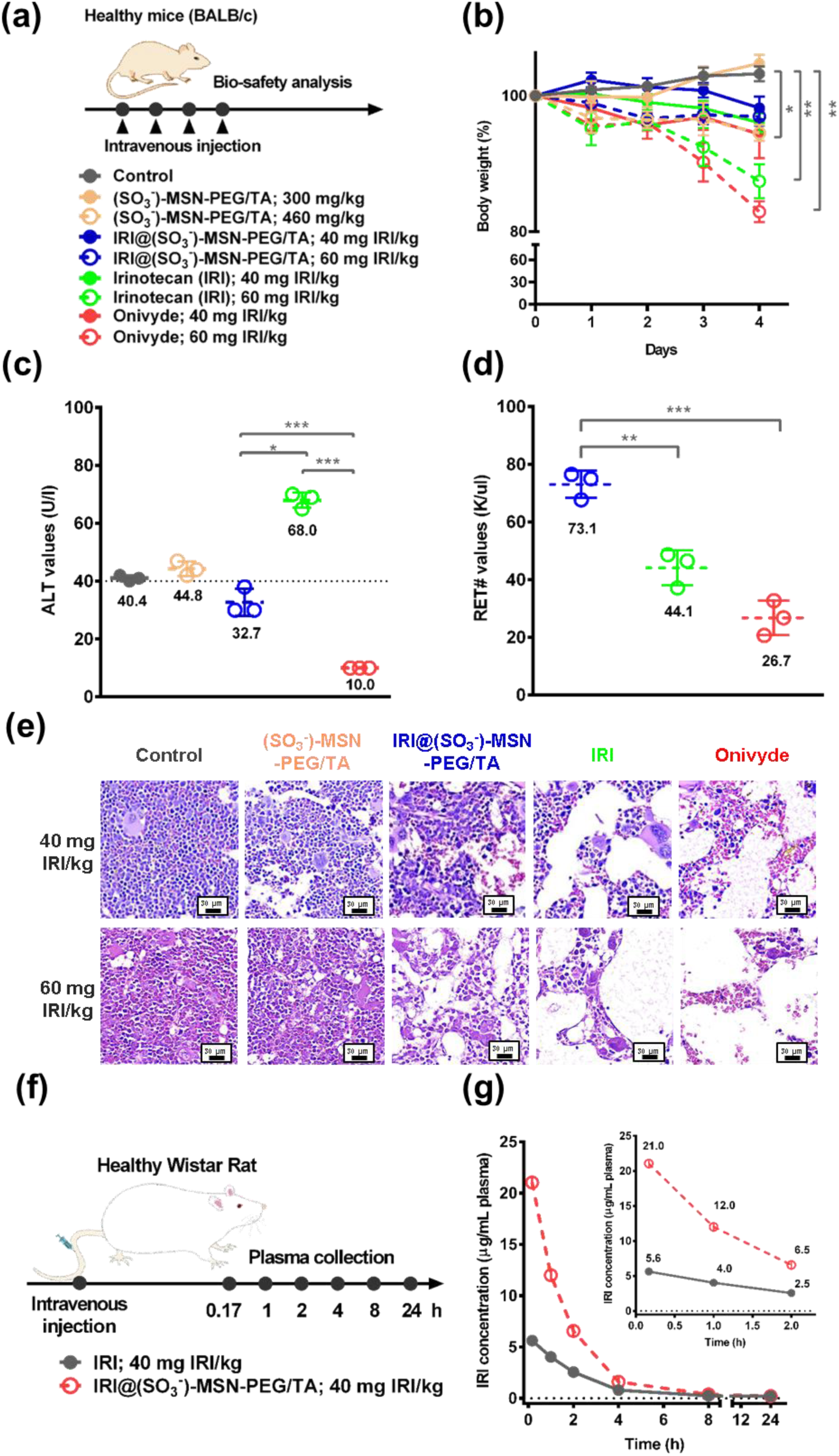
In vivo biosafety and pharmacokinetic analysis of IRI@(SO_3_^-^)-MSN-PEG/TA in healthy animal model. (a) Schematic representation of the multiple-dose toxicity study. Healthy BALB/c mice were intravenously injected with (SO_3_^-^)-MSN-PEG/TA (300 and 460 mg/kg), free IRI (40 and 60 mg IRI/kg), IRI@(SO_3_^-^)-MSN-PEG/TA (equivalent IRI dose), or Onivyde (equivalent IRI dose) once daily for four consecutive administrations. (b) Body weight variations across different treatment groups. (*p < 0.05, ** p < 0.01, n = 4) (c) Serum alanine transaminase (ALT) levels after four administrations of (SO_3_^-^)-MSN-PEG/TA (460 mg/kg), IRI, IRI@(SO_3_^-^)-MSN-PEG/TA, or Onivyde (each 60 mg IRI/kg) (*p < 0.05, ***p < 0.001, n = 4). (d) Reticulocyte (RET#) counts from CBC analysis following four administrations of IRI@(SO_3_^-^)-MSN-PEG/TA, IRI, or Onivyde (60 mg IRI/kg) (**p < 0.01, ***p < 0.001, n = 4). (e) Representative histopathological analysis of bone marrow from healthy BALB/c mice. Hematoxylin and eosin (H&E)-stained sections were obtained from mice treated with IRI alone (40 and 60 mg IRI/kg), IRI@(SO_3_^-^)-MSN-PEG/TA, (SO_3_^-^)-MSN-PEG/TA, or Onivyde (equivalent IRI dose) once daily for four administrations. Scale bar = 30 µm. (f) Pharmacokinetic study design and (g) plasma concentration–time profiles. Healthy Wistar rats received a single intravenous injection of free IRI (40 mg/kg) or IRI@(SO_3_^-^)-MSN-PEG/TA (40 mg IRI/kg, N = 4). Plasma concentrations of IRI were quantified by LC-MS/MS at 0.17, 1, 2, 4, 8, and 24 hours’ post-injection. Insets highlight early-phase IRI kinetics (0.17–2 hours).

To further evaluate systemic safety, complete blood count (CBC) and serum biochemical analyses were performed (Figure 5c–d, Tables S4–S5). Serum biochemistry showed that alanine transaminase (ALT) levels were abnormally low in Onivyde-treated mice (10 U/L), possibly reflecting liver dysfunction and metabolic frailty (Figure 5c and Table S4, red-labeled).[47, 48] In contrast, ALT levels in the IRI@(SO_3_^-^)-MSN-PEG/TA group (32.7 U/L) remained within the normal range (28–132 U//L)[49, 50] Although ALT levels in the high-dose IRI group (68.0 U/L) were also within the normal range, they were elevated relative to controls (40.4 U/L), suggesting mild hepatic damage likely resulting in increased ALT release into the serum.. CBC analysis revealed that high-dose IRI and Onivyde treatments (60 mg IRI/kg) caused a marked reduction in reticulocyte counts (RET#, IRI: 44.1 K/μL; Onivyde: 26.7 K/μL) compared with controls (498 K/μL), indicating impaired erythropoiesis (Figure 5d and Table S5, red-labeled). Notably, IRI@(SO_3_^-^)-MSN-PEG/TA (60 mg IRI/kg) treatment partially restored reticulocyte levels (73.1 K/μL), suggesting that nanoencapsulation alleviated hematopoietic toxicity.

Histopathological analysis of bone marrow (Figure 5e) further supported these findings. Mice treated with free IRI or Onivyde (40 and 60 mg IRI/kg) exhibited pronounced myelosuppression, characterized by decreased hematopoietic cellularity and reduced bone-marrow density.[12, 51] In contrast, IRI@(SO_3_^-^)-MSN-PEG/TA and (SO_3_^-^)-MSN-PEG/TA groups maintained relatively preserved marrow cellularity, indicating that encapsulating IRI in (SO_3_^-^)-MSN-PEG/TA substantial mitigatied bone marrow toxicity. As expected, all 60 mg IRI/kg groups displayed greater marrow suppression than their 40 mg IRI/kg counterparts. Liver and intestinal tissues were also examined in the high-dose (60 mg IRI/kg) treatment groups. Mild mononuclear inflammatory infiltration was observed in the liver across all IRI-containing groups, but no treatment-specific necrosis or architectural disruption was detected. Similarly, intestinal sections exhibited intact crypt architecture without epithelial necrosis (Figure S3). These observations collectively underscore the dual advantages of IRI@(SO_3_^-^)-MSN-PEG/TA, enhanced antitumor efficacy coupled with markedly reduced systemic and hematopoietic toxicity, highlighting (SO_3_^-^)-MSN-PEG/TA as a promising nanocarrier to reduce the adverse effects of IRI-based therapy.

Based on findings related to systemic toxicity, a dose of 40 mg IRI/kg was chosen for pharmacokinetic (PK) evaluation. Healthy Wistar rats received a single intravenous injection of either free IRI or IRI@(SO_3_^-^)-MSN-PEG/TA, and plasma samples were collected at predetermined time points up to 24 hours (0.17, 1, 2, 4, 8, and 24 hours) (see Figure 5f). As shown in Figure 5g and Table 1, IRI@(SO_3_^-^)-MSN-PEG/TA exhibited a substantially higher plasma concentration compared with free irinotecan. The maximum concentration (C_max_) of IRI@(SO_3_^-^)-MSN-PEG/TA reached 21.0 µg/mL, representing a 3.7-fold increase over free IRI (5.6 µg/mL). Systemic exposure, as measured by the area under the curve (AUC_0-∞_), was 45.1 h·µg/mL for IRI@(SO_3_^-^)-MSN-PEG/TA, showing a 2.4-fold increase over free IRI (18.8 h·µg/mL). In addition to enhanced exposure, IRI@(SO_3_^-^)-MSN-PEG/TA exhibited a slower clearance rate (CL) and decreased volume of distribution (Vd). While MSN formulation improved systemic exposure (C_max_ and AUC_0-∞_) with reduced CL and Vd, the apparent half-life (t_1/2_) of IRI@(SO_3_^-^)-MSN-PEG/TA was slightly shorter than that of free IRI. This discrepancy may be attributed to the sampling schedule; for free IRI, the first blood collection at 10 minutes’ post-injection likely missed the rapid alpha-phase distribution, resulting in a slightly longer calculated t_1/2_. Given that free IRI rapidly distributes into tissues and is cleared more quickly (higher Vd and CL), as well as undergoes rapid conversion from the active lactone form to the inactive carboxylate form under physiological conditions, the actual systemic half-life of free IRI is expected to be shorter than the observed value.[52]

**Table 1.** Pharmacokinetic Parameters of IRI and IRI@(SO_3_^-^)-MSN-PEG/TA.

| Parameters | Unit | IRI | IRI@ (SO<sub>3</sub><sup>-</sup>)-MSN-PEG/TA |
| --- | --- | --- | --- |
| dose | mg/kg | 40 | 40 |
| C<sub>max</sub> | μg/mL | 5.6 | 21.0 |
| AUC<sub>0–∞</sub> | $h \cdot \mu$g/mL | 18.8 | 45.1 |
| T<sub>1/2</sub> | h | 5.3 | 4.0 |
| CL | L/h/kg | 2.1 | 0.9 |
| Vd | L/kg | 16.3 | 5.0 |

C<sub>max</sub>, maximum observed concentration; AUC<sub>0– ∞</sub> , area under the plasma concentration–time curve from time zero to infinity; T<sub>1/2</sub>, apparent half-life; CL, clearance; Vd, volume of distribution.

The PK profile of IRI@(SO_3_^-^)-MSN-PEG/TA demonstrates improved systemic exposure and provides a mechanistic basis for its superior antitumor efficacy and reduced systemic toxicity compared with free IRI. Importantly, the therapeutic design of IRI-loaded MSNs enables rapid tumor targeting and timely clearance. The drug-release half-time of approximately 5 hours (Figure 1i) allows sufficient circulation to achieve tumor accumulation via the EPR effect (typically within 1 h). However, its shorter blood circulation time also helps minimize systemic toxicity, particularly in sensitive tissues such as the bone marrow. In contrast, lipid-based nanoparticles such as silicasomes and Onivyde exhibit extended circulation (several days), promoting gradual tumor accumulation but increasing the risk of off-target deposition in sensitive tissues, leading to potential toxicities.

Additionally, IRI is activated by carboxylesterase 2 (CE2), which has two forms, the more effective one in Endoplasmic reticulum (ER)-retained form and the less active form of secreted CE2[53]. MSNs enter cells via clathrin-mediated endocytosis[54, 55] and localize near the ER, enhancing CE2-mediated conversion of IRI to SN-38. In contrast, Onivyde enters cells mainly through plasma membrane fusion, with limited ER access and thus activate IRI less. At the same time, IRI released near cell membrane leads to rapid P-gp efflux, thus to drug resistance.[56–58]

## Conclusion

In conclusion, this study introduces a surface functionalized MSNs ((SO_3_^-^)-MSN-PEG/TA) as a dual-functional drug delivery for mCRC therapy, cytotoxic and anti-metastasis. The unique design of (SO_3_^-^)-MSN-PEG/TA not only stabilizes IRI by preserving its active lactone form at over 99% for up to six months, but also enhances drug loading efficiency and enables sustained release, successfully overcoming long-standing bottlenecks and limitations of IRI treatment. Moreover, (SO_3_^-^)-MSN-PEG/TA exhibits intrinsic antimetastatic properties by modulating focal adhesion dynamics, effectively inhibiting tumor migration and angiogenesis, while achieving efficient tumor accumulation through the EPR effect. Both in vitro and in vivo results confirmed that IRI@(SO_3_^-^)-MSN-PEG/TA provides superior antitumor efficacy and antimetastatic potential, significantly suppressing primary tumor growth and metastatic spread. Importantly, pharmacokinetic analysis revealed improved systemic retention and bioavailability of IRI compared with free drug, supporting enhanced therapeutic durability. This nanoformulation also demonstrated excellent biosafety, preserving bone marrow cellularity and mitigating adverse effects typically associated with free IRI and Onivyde. By integrating potent chemotherapy with intrinsic metastasis suppression and favorable pharmacokinetics, IRI@(SO_3_^-^)-MSN-PEG/TA represents a significant advancement and a safer, more effective therapeutic strategy for mCRC. Future efforts should focus on advancing this platform toward clinical translation and exploring its applicability across other metastatic cancers.

## Acknowledgments

The authors gratefully acknowledge the Laboratory Animal Center at Taipei Medical University (TMU) for technical support in animal experiments. The TOC graphic was created using BioRender.com. This research was supported by Taiwan’s National Science and Technology Council (NSTC112-2113-M-038-004, NSTC112-2113-M-038-002, NSTC113-2113-M-001-033, and NSTC113-2124-M-038-002), the TMU Industry-Academic Cooperation Program (A-105-074, A-109-121, and A-113-077), and Academia Sinica of Taiwan (AS-IA-110-M04 and AS-GCS-113-M01).

## TOC

IRI@(SO_3_^-^)-MSN-PEG/TA highlights its ability to preserve the active lactone form of IRI for up to six months, suppress metastasis, enhance tumor accumulation via the EPR effect, minimize systemic toxicity, and significantly improve overall survival, representing a safe and effective nanomedicine for metastatic colorectal cancer treatment.

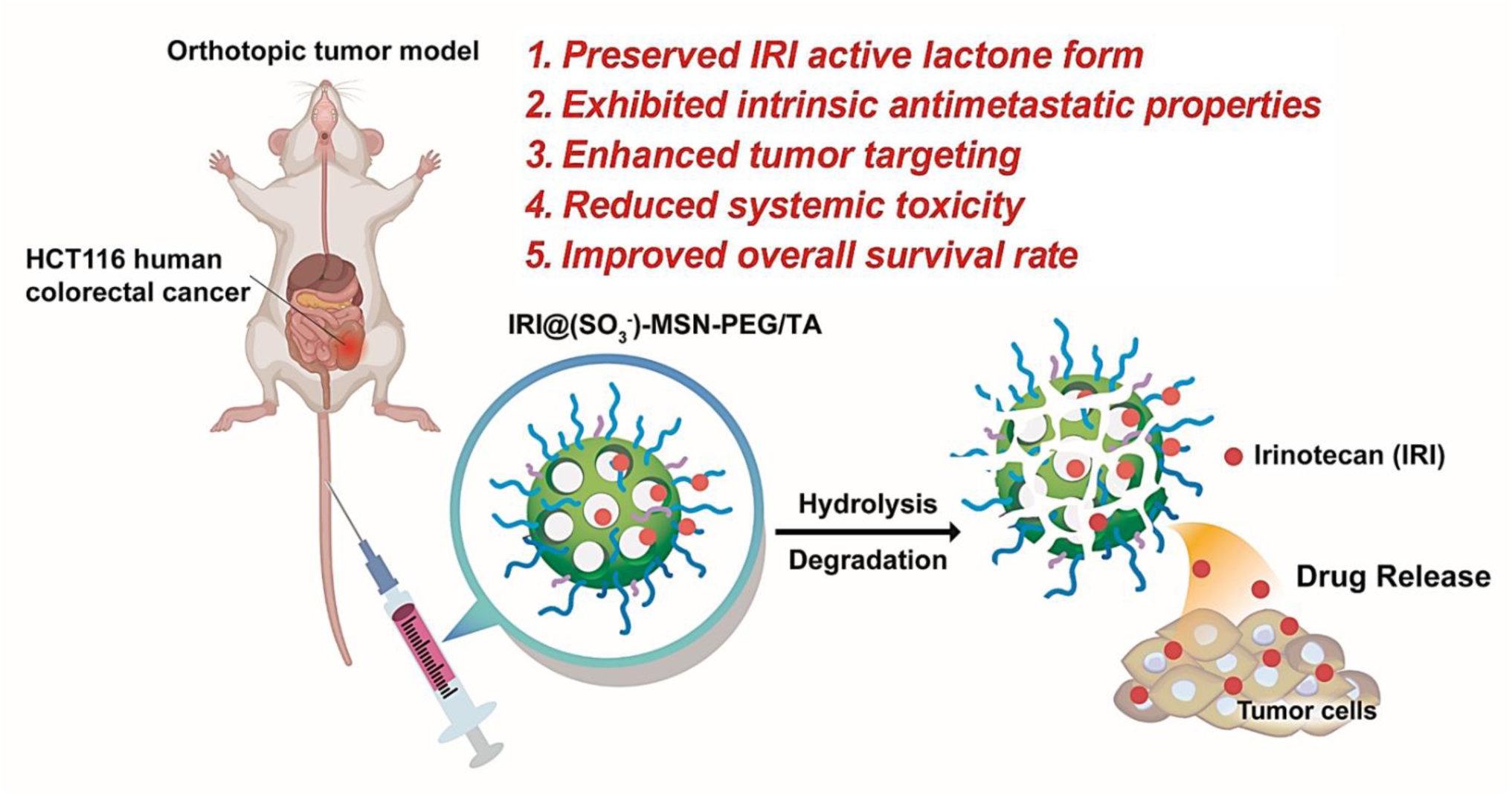

