## Supporting Information for "Antimetastatic Sulfonate-Functionalized Mesoporous Silica Nanoparticles Enhance Irinotecan Stability and Delivery for Colorectal Cancer Treatment"

**Table S1.** Characteristics of functionalized MSNs and IRI encapsulation

| **Samples** | **Z-average in PBS (d.nm) / PdI** | **Zeta potential (mV)** | **TEM size (nm)** | **LC%** | **LE%** |
|---|---|---|---|---|---|
| MSN-PEG/TA | 40.3 / 0.07 | -8.1 ± 0.5 | 28.5 ± 4.5 | | |
| ($SO_3^-$)-MSN-PEG/TA | 38.7 / 0.07 | -28.2 ± 0.8 | 25.5 ± 4.0 | | |
| IRI@MSN-PEG/TA | 40.1 / 0.08 | | | 3.9 ± 0.5% | 45 ± 7% |
| IRI@($SO_3^-$)-MSN-PEG/TA | 39.9 / 0.09 | | | 11.5 ± 1.0% | 95 ± 4% |

Z-average = harmonic intensity averaged particle diameter, PDI = polydispersity index. IRI = Irinotecan. Zeta potential, representing the surface charge of MSNs, was measured in diluted PBS solution. LC%, IRI loading capacity; LE%, IRI loading efficiency

**Table S2.** Elemental analysis of functionalized MSNs

| | **Weight for analysis (mg)** | **Elemental analysis** | | | |
|---|---|---|---|---|---|
| | | **C%** | **N%** | **S%** | **H%** |
| MSN-PEG/TA | 3.0 | 15.8 ± 0.1 | 0.10 ± 0.01 | ND | 4.22 ± 0.2 |
| ($SO_3^-$)-MSN-PEG/TA | 2.7 | 21.0 ± 0.2 | 0.07 ± 0.01 | 2.25 ± 0.1 | 5.22 ± 0.1 |

Percentages of carbon, nitrogen, sulfur and hydrogen contents for functionalized MSNs were obtained from an elemental analysis. ND, not detected

**Table S3** Thermogravimetric analysis (TGA) functionalized MSNs

| Samples | TGA results for 40°C-200°C (wt%) | TGA results for 200°C-600°C (wt%) | TGA results for 600°C-800°C (wt%) |
|---|---|---|---|
| MSN | 5.6% | 2.4% | 0.8% |
| MSN-PEG | 6.2% | 24.9% | 0.7% |
| MSN-PEG-TA | 1.5% | 33.9% | 0.9% |
| ($SO_3^-$)-MSN-PEG-TA | 1.7% | 37.5% | 1.0% |

wt%: normalized weight loss from thermogravimetric analysis (TGA) analysis.

**Table S4.** Biochemical parameters in healthy BALB/c mice after various treatments.

| Test description | Unit | Control | $(SO_3^-)$-MSN-PEG/TA | | IRI@$(SO_3^-)$-MSN-PEG/TA | | IRI | | Onivyde | |
|---|---|---|---|---|---|---|---|---|---|---|
| | | | 300 mg/kg | 460 mg/kg | 40 mg IRI/kg | 60 mg IRI/kg | 40 mg IRI/kg | 60 mg IRI/kg | 40 mg IRI/kg | 60 mg IRI/kg |
| BUN | (mg/dL) | 22.5 ± 1.3 | 27 ± 5.56 | 31 ± 4.2 | 21 ± 3.6 | 20.2 ± 6.1 | 22.3 ± 3.5 | 22.25 ± 3.7 | 19.7 ± 3.2 | 18 ± 2.9 |
| CREA | (mg/dL) | 0.1 ± 0 | 0.1 ± 1.7 | 0.1 ± 0 | 0.1 ± 0 | 0.1 ± 0 | 0.1 ± 0 | 0.1 ± 0 | 0.2 ± 0.17 | 0.1 ± 0 |
| TP | (g/dL) | 5.25 ± 0.2 | 5 ± 0.2 | 5.1 ± 0.1 | 5.2 ± 0.2 | 4.98 ± 0.1 | 5.3 ± 0.2 | 5.325 ± 0.5 | 4.9 ± 0.8 | 5.4 ± 0.08 |
| ALB | (g/dL) | 2.475 ± 0.2 | 2.3 ± 0.1 | 2.3 ± 0.1 | 2.5 ± 0.1 | 2.26 ± 0.1 | 2.5 ± 0.1 | 2.5 ± 0.2 | 2.2 ± 0.5 | 2.775 ± 0.2 |
| ALT | (U/L) | 40.4 ± 1.5 | 45 ± 6.1 | 44.8 ± 2.2 | 45.3 ± 4 | 32.7 ± 4.6 | 41.3 ± 2.6 | 68.0 ± 2.6 | 10 ± 0.03 | 10 ± 0.05 |
| AST | (U/L) | 85.5 ± 14.2 | 83.7 ± 12.6 | 80.5 ± 6.8 | 80.7 ± 10.6 | 111.6 ± 18 | 145.7 ± 40 | 128.3 ± 40 | 89 ± 21 | 122.8 ± 29 |
| ALKP | (U/L) | 122.8 ± 20 | 158.7 ± 38 | 135.5 ± 28 | 120 ± 15.5 | 122.8 ± 6.9 | 140.3 ± 13 | 104.8 ± 35 | 147.3 ± 27 | 134.3 ± 13 |
| LDH | (U/L) | 1085 ± 187 | 1253 ± 554 | 1083 ± 131 | 685 ± 267 | 1256 ± 394 | 1992 ± 478 | 2697 ± 681 | 1463 ± 243 | 1024 ± 183 |

BUN, blood urea nitrogen; CREA, creatinine; TP, total protein; ALB, albumin; ALT, alanine transaminase; AST, aspartate transaminase; ALKP, alkaline phosphatase, LDH, lactate dehydrogenase.

**Table S5.** Hematological parameters of the complete blood count in healthy BALB/c mice after various treatments.

| Test description | Unit | Control | ($SO_3^-$)-MSN-PEG/TA | | IRI@($SO_3^-$)-MSN-PEG/TA | | IRI | | Onivyde | |
|---|---|---|---|---|---|---|---|---|---|---|
| | | | 300 mg/kg | 460 mg/kg | 40 mg IRI/kg | 60 mg IRI/kg | 40 mg IRI/kg | 60 mg IRI/kg | 40 mg IRI/kg | 60 mg IRI/kg |
| RBC | (M/μL) | 11.1 ± 0.7 | 11.4 ± 0.5 | 11 ± 0.9 | 10.4 ± 0.2 | 11 ± 0.3 | 10.2 ± 0.3 | 10.2 ± 0.6 | 11.6 ± 0.2 | 11.2 ± 0.2 |
| HGB | (g/dL) | 16.1 ± 0.9 | 16.9 ± 0.9 | 16.4 ± 1.1 | 15.3 ± 0.1 | 16.1 ± 0.3 | 14.8 ± 0.6 | 15 ± 0.9 | 17.1 ± 0.2 | 16.6 ± 0.4 |
| HCT | (%) | 54.4 ± 3.6 | 57.2 ± 3 | 54.4 ± 5.1 | 49.7 ± 1.4 | 52.2 ± 1.1 | 48.4 ± 2.2 | 48.6 ± 3.4 | 55.9 ± 0.7 | 53.1 ± 1.7 |
| MCV | (fL) | 49.2 ± 0.3 | 50.3 ± 0.3 | 49.7 ± 0.8 | 47.6 ± 0.6 | 47.7 ± 0.6 | 47.3 ± 0.7 | 47.6 ± 0.3 | 48.4 ± 0.6 | 47.3 ± 0.7 |
| MCH | (pg) | 14.5 ± 0.1 | 14.9 ± 0.1 | 15 ± 0.3 | 14.7 ± 0.2 | 14.7 ± 0.1 | 14.5 ± 0.2 | 14.7 ± 0.1 | 14.8 ± 0.1 | 14.8 ± 0.1 |
| MCHC | (g/dL) | 29.6 ± 0.3 | 29.6 ± 0.2 | 30.2 ± 0.9 | 30.8 ± 0.7 | 30.9 ± 0.2 | 30.6 ± 0.3 | 30.8 ± 0.4 | 30.5 ± 0.3 | 31.3 ± 0.6 |
| RDW-SD | (fL) | 31.4 ± 2.5 | 31.3 ± 0.9 | 32 ± 1.8 | 29.1 ± 1.2 | 28.6 ± 1.5 | 29.4 ± 1 | 28 ± 0.4 | 32.3 ± 1.3 | 29.6 ± 2.2 |
| RDW-CV | (%) | 25.8 ± 1.6 | 26 ± 0.9 | 26.3 ± 1.4 | 24.6 ± 0.3 | 24.9 ± 0.6 | 24.8 ± 0.6 | 23.6 ± 0.5 | 27.2 ± 1.1 | 25.6 ± 1 |
| RET# | (K/μL) | 498.3 ± 77 | 460.2±81 | 470 ± 73 | 90.9 ± 12 | 73.1 ± 4.7 | 95.3 ± 12 | 44.1 ± 6.1 | 47.4 ± 5.8 | 26.7 ± 6 |
| RET% | (%) | 4.5 ± 0.5 | 3.9 ± 1.4 | 6.3 ± 1.7 | 0.9 ± 0.1 | 0.7 ± 0 | 0.9 ± 0.1 | 0.5 ± 0.1 | 0.4 ± 0.1 | 0.3 ± 0.1 |
| PLT | (K/μL) | 788 ± 52.1 | 704.3±341 | 452.9 ± 64 | 615 ± 95.6 | 380.7±89 | 765.3±111 | 665 ± 17.3 | 713.8 ± 37 | 685.6 ± 51 |
| WBC | (K/μL) | 5 ± 2.4 | 6.7 ± 1.7 | 5.1 ± 2.3 | 3.7 ± 0.5 | 2.3 ± 1.2 | 3.5 ± 1.3 | 1.3 ± 0 | 6.8 ± 0.7 | 2.3 ± 1.4 |
| NEUT# | (K/μL) | 1.4 ± 0.8 | 1.7 ± 0.4 | 1.6 ± 0.6 | 1.1 ± 0.2 | 0.5 ± 0.2 | 1 ± 0.8 | 0.4 ± 0.2 | 1.4 ± 0.5 | 0.5 ± 0.3 |
| LYMPH# | (K/μL) | 3.4 ± 1.7 | 4.7 ± 1.3 | 3.3 ± 1.6 | 2.6 ± 0.2 | 1.7 ± 1.1 | 2.4 ± 0.7 | 0.9 ± 0.2 | 5.3 ± 0.3 | 1.7 ± 1.1 |
| MONO# | (K/μL) | 0.1 ± 0 | 0.1 ± 0 | 0.1 ± 0.1 | 0.1 ± 0 | 0 ± 0 | 0.1 ± 0 | 0 ± 0 | 0.1 ± 0 | 0 ± 0 |
| EO# | (K/μL) | 0.1 ± 0 | 0.2 ± 0 | 0.1 ± 0 | 0 ± 0 | 0 ± 0 | 0 ± 0 | 0 ± 0 | 0 ± 0 | 0 ± 0 |
| BASO# | (K/μL) | 0 ± 0 | 0 ± 0 | 0 ± 0 | 0 ± 0 | 0 ± 0 | 0 ± 0 | 0 ± 0 | 0 ± 0 | 0 ± 0 |

| | | | | | | | | | | |
|---|---|---|---|---|---|---|---|---|---|---|
| NEUT% | (%) | 28.9 ± 6.1 | 25.1 ± 1.1 | 31.9 ± 4.9 | 29 ± 2.6 | 26.1 ± 14 | 24.8 ± 16 | 28.1 ± 16 | 20.2 ± 5.1 | 23.5 ± 5 |
| LYMPH% | (%) | 68.4 ± 6.5 | 70.6 ± 1.8 | 63.4 ± 4.6 | 68.3 ± 2.7 | 71.7 ± 15 | 72.3 ± 15 | 70.9 ± 16 | 78.3 ± 5.4 | 74.9 ± 5.2 |
| MONO% | (%) | 1 ± 0.5 | 1.9 ± 0.2 | 2.2 ± 0.5 | 2 ± 0.3 | 1.5 ± 0.3 | 2.6 ± 1.1 | 1 ± 0.4 | 1.3 ± 0.5 | 1.3 ± 0.5 |
| EO% | (%) | 1.5 ± 0.1 | 2.4 ± 1 | 2.1 ± 0.4 | 0.6 ± 0.4 | 0.5 ± 0.8 | 0.2 ± 0.2 | 0 ± 0 | 0.1 ± 0.1 | 0.1 ± 0.2 |
| BASO% | (%) | 0.3 ± 0.1 | 0.1 ± 0.1 | 0.4 ± 0.3 | 0.1 ± 0.1 | 0.3 ± 0.5 | 0.2 ± 0.3 | 0 ± 0 | 0.1 ± 0.1 | 0.2 ± 0.3 |

Abbreviations: RBC, red blood cells; HGB, hemoglobin; HCT, hematocrit; MCV, mean corpuscular volume; MCH, mean corpuscular hemoglobin; MCHC, mean corpuscular hemoglobin concentration; RDW-SD, red blood cell distribution width standard deviation; RDW-CV, red blood cell distribution width expressed as a coefficient of variation; RET, reticulocytes; PLT, platelet count; NEUT, neutrophil count; LYMPH, lymphocyte count; MONO, mononucleosis; EO, eosinophils; BASO, basophils.

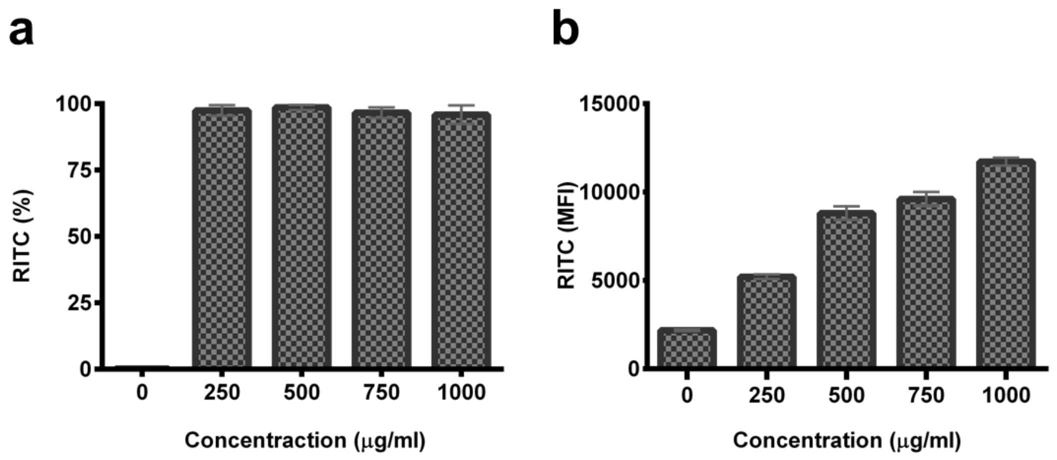


**Figure S1.** The cellular uptake of RITC conjugated ($SO_3^-$)-MSN-PEG/TA at different concentrations in HCT-116 cells was measured by flow cytometry after 24 hours of treatment. (a) Percentage and (b) mean fluorescence intensity (MFI) of positive cells.

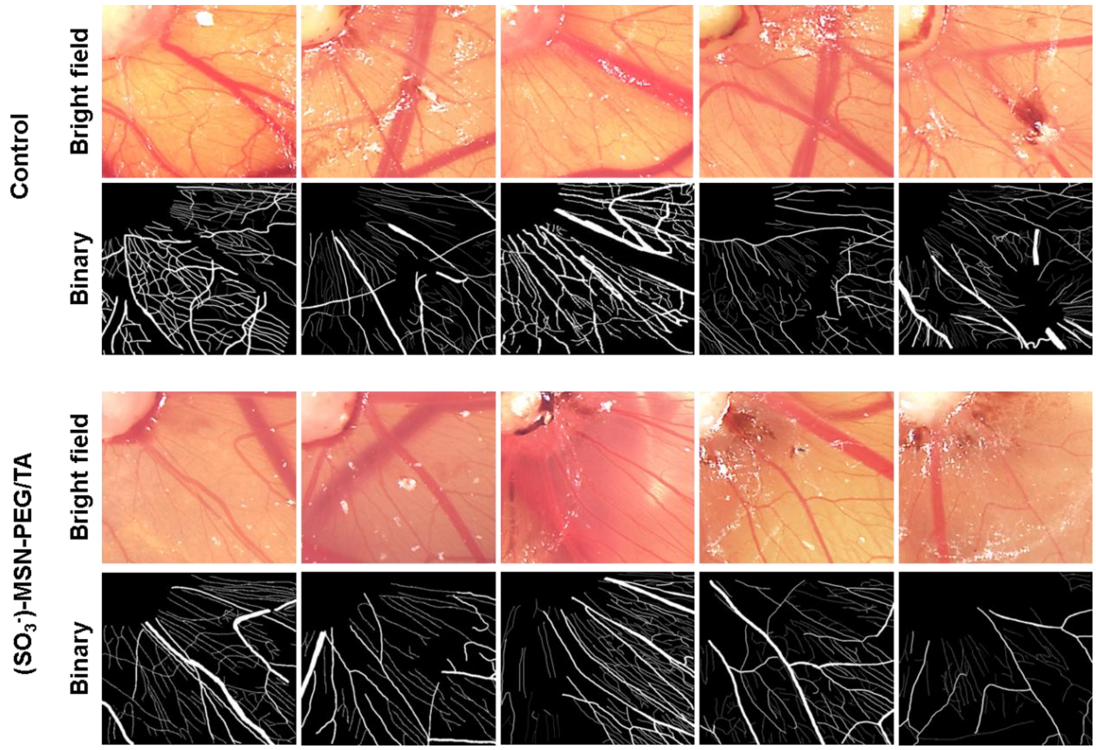


**Figure S2.** The quantitative analysis of vascular density in the chick CAM. Blood vessel densities were calculated by NIH ImageJ software with the "angiogenesis analyzer" plug-in.

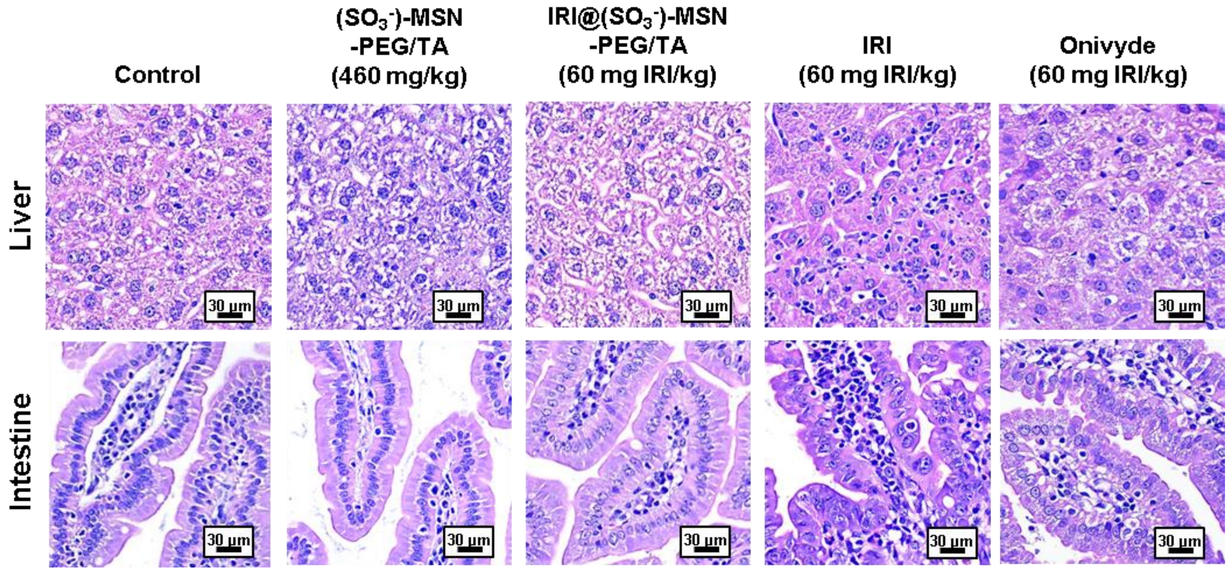


**Figure S3.** Histopathological analysis of liver and intestine from healthy BALB/c mice. H&E-stained sections were obtained from mice treated with IRI (60 mg IRI/kg), IRI@($SO_3^-$)-MSN-PEG/TA, ($SO_3^-$)-MSN-PEG/TA, or Onivyde (equivalent to IRI dose) daily for a total of four administrations. Scale bar = 30 μm.
